# Phytoplasma mediated transcriptional changes in poinsettia buds suggest MAF3 and bZIP67 transcription factors as potential suppressors of shoot branching

**DOI:** 10.64898/2026.06.16.732632

**Authors:** Behrooz Darbani, Christina R. Ingvardsen, Inger B. Holme, Marianne Gellert Møller, Jacob Graff, Henrik Brinch-Pedersen, Mogens Nicolaisen

## Abstract

Shoot branching is critical not only in breeding for yield but also for ornamentals’ architecture. In the ornamental plant poinsettia (*Euphorbia pulcherrima*), shoot branching has traditionally been induced by phytoplasma (*Candidatus* Phytoplasma pruni) inoculation. This study aimed to identify regulatory genes that could be leveraged in future breeding-by–genetic engineering efforts to develop phytoplasma-free, branching poinsettia plants.
To elucidate mechanisms of phytoplasma-induced shoot branching, we performed RNA-sequencing and assembled an axillary bud-specific transcriptome for expression analyses in phytoplasma-infected and -free poinsettia. Phenotyping and RNA-sequencing were also conducted on *Arabidopsis* mutants and wild-type lines to investigate the transcriptional regulatory effects of candidate genes.
The transcription factors *Ep*MAF3 and *Ep*bZIP67 were highly de-regulated in phytoplasma-infected poinsettia. We also found a two-fold increase in primary-stem branching levels of the *Arabidopsis maf*3 and *bzip*67 mutants, suggesting the two transcription factors as potential shoot branching suppressors. *AtTcp*1, a CYC-clade TCP transcription factor, was up-regulated (78x) in leaves of the *maf*3 mutants. Analyzing previously reported protein-level interactions for the differentially expressed genes (e.g., *AtClamt, AtGh*3.9/3.15, *AtSaur*32/36, *AtAbi*3, *AtGamt*2, *AtTcp*3, and *AtDwf*4) in *bzip*67 mutants shed light on other shoot branching regulators such as TCPs, PINs, ABIs, DWARF14, and BES1, highlighting two regulatory sub-networks including membrane transport and hormonal signaling.
The results open the way to rational engineering of shoot branching in poinsettia by targeted mutagenesis of MAF3 and bZIP67. In that way, the tedious and viral-infection prone process of phytoplasma inoculation can be avoided and poinsettia plants would have more homogenous branching.

**One-sentence summary:** Phytoplasma infection in poinsettia induces bud-specific repression of the transcription factors *EpMaf*3 and *EpbZip*67, consistent with the enhanced stem branching observed in *Arabidopsis maf*3 and *bzip*67 mutant lines.

## Introduction

The level of stem branching in plants affect crop yields, but also the marketability of ornamentals. Shoot branching involves axillary meristem development and the growth of axillary buds. However, the outgrowth of axillary buds is inhibited by the apical meristem, a process known as apical dominance, at different levels depending on the plant species. The inhibition of axillary buds involves hormonal regulation, such as auxins, cytokinins, abscisic acid, and strigolactones leading to downstream adjustments such as sugar starvation in axillary buds (Wang *et al*. 2018; Beveridge *et al*. 2023). The gene Teosinte Branched1, also known as *Brc*1, is one of the central actors regulating branching (Doebley *et al*. 1995, 1997; Aguilar-Martínez *et al*. 2007). In *Arabidopsis thaliana*, BRANCHED1 (BRC1/TCP18) and BRANCHED2 (BRC2/TCP12), as the closest homologues for rice Teosinte Branched1, repress stem branching (Aguilar-Martínez *et al*. 2007). These are members of the group of TCP transcription factors involved in plant growth and developmental regulation (Martín-Trillo & Cubas 2010; Nicolas & Cubas 2016). Rice Ideal Plant Architecture1 (IPA1) has been found to positively regulate Teosinte Branched1 by binding to its promoter (Lu *et al*. 2013). Interestingly, the promoter regions of up- and downregulated genes after decapitation, which promotes axillary bud growth, are enriched in a sugar response motif and TCP binding sites, respectively (Tatematsu *et al*. 2005). While sugars and cytokinin inhibit the BRC1 mediated growth suppression of axillary buds, auxins and strigolactones promote this process (Wang *et al*. 2018).

Strigolactone biosynthesis genes such as *Dwarf*27 (Lin *et al*. 2009), *Max*1 (Stirnberg *et al*. 2002; Booker *et al*. 2005), *Max*3 (Booker *et al*. 2004, 2005; Schwartz *et al*. 2004), *Max*4 (Sorefan *et al*. 2003; Schwartz *et al*. 2004; Booker *et al*. 2005), and strigolactone signaling genes such as *Max*2 (Stirnberg *et al*. 2002, 2007; Booker *et al*. 2005) and *Dwarf*14 (Hamiaux *et al*. 2012; Nakamura *et al*. 2013 14) are involved in negative regulation of the suppression of BRC1. This is facilitated by MAX2 and DWARF14 mediated degradation of BRC1 suppressors *At*BES1/*Os*BZR1 and *At*SMXLs/*Os*D53 (Wang *et al*. 2013; Hu *et al*. 2019; Fang *et al*. 2020 1). The identification of *AtBes*1/*OsBzr*1 as the strigolactone signaling targets reveals the involvement of brassinosteroids in shoot branching regulation; BES1 and BZR1 are stabilized and accumulate within the nucleus in response to brassinosteroids (Yin *et al*. 2002). While auxin signaling positively regulates strigolactone biosynthesis (Hayward *et al*. 2009) and thereby the expression of *Brc*1, BRC1 also represses the expression of the auxin efflux carrier gene *Pin*3 and thereby the accumulation of auxin in axillary buds (Shen *et al*. 2019). Likewise, BRC1 contributes to the suppression of branching via abscisic acid accumulation as a result of the positive transcriptional regulation of HOMEOBOX 21 (*Hb*21), HOMEOBOX 40 (*Hb*40), and HOMEOBOX 53 (*Hb*53) (González-Grandío *et al*. 2017). In contrast, the wheat ankyrin repeat protein coding gene Tiller Number1 (TN1) suppresses abscisic acid biosynthesis via interaction with abscisic acid receptors called pyrabactin resistance 1 like proteins (PYLs) and thereby increases tillering (Dong *et al*. 2023). While auxins, strigolactones and abscisic acid suppress axillary bud growth, cytokinins and sucrose enhances bud outgrowth. Sucrose promotes stem branching through cytokinin biosynthesis (Salam *et al*. 2021). It has also been shown that sucrose may interfere with strigolactone signaling and *Brc*1 expression and consequently trigger axillary bud outgrowth and tillering via transcriptional suppression of *Max*2/D3 which in turn leads to the accumulation of D53 (Barbier *et al*. 2015; Patil *et al*. 2022). Furthermore, the expression of trehalose 6-phosphate phosphatase lowers trehalose 6-phosphate levels in axillary buds and consequently inhibits branching (Fichtner *et al*. 2021). Like the rice IPA1, which is a SQUAMOSA PROMOTER BINDING PROTEIN-LIKE (SPL) transcription factor and positively regulates the expression of Teosinte Branched1 (Lu *et al*. 2013), tomato *Spl*13 is transcriptionally upregulated by strigolactones and inhibits cytokinin biosynthesis resulting in activation of *Brc*1 transcription (Chen *et al*. 2023). As the case with cytokinins, bioactive gibberellins such as GA1 and GA4 promote branching through interfering with strigolactone signaling (Ni *et al*. 2015; Katyayini *et al*. 2020). In contrast, gibberellins such as GA3 and GA6 have interfering effects on branching (Katyayini *et al*. 2020). These bioactive forms of gibberellins are biosynthesized by the action of GA3-oxidase (*G3ox*) genes and can be converted to inactive forms by *G2ox* genes (Rieu *et al*. 2008; Katyayini *et al*. 2020).

Phytoplasmas, as plant obligate parasites, code for secreted effector proteins that interact with different plant signaling proteins intracellularly leading to plant dwarfism, increased branching, and bushy structures (Hoshi *et al*. 2009). Phytoplasmas code for ≈ 40-60 effector proteins which are, via the Sec system, secreted into the cytoplasm of the host (Bai *et al*. 2009; Oshima *et al*. 2023). For example, the phytoplasma SAP11 effector protein destabilizes TCP transcription factors such as BRC1 resulting in a dwarfed and bushy architecture (Sugio *et al*. 2011; Chang *et al*. 2018; Pecher *et al*. 2019). SAP54 is another effector protein reported to mediate the degradation of different MADS-box transcription factors of the host (MacLean *et al*. 2014; Kitazawa *et al*. 2022). In addition to the developmental effects, phytoplasma effector proteins also interact with host defense mechanisms, for example through the interaction of the effector protein SWP12 with wheat WRKY74 (Bai *et al*. 2023), to increase the host susceptibility to phytoplasma.

As one of the popular ornamental potted plants, poinsettia (*E. pulcherrima*; known as Christmas Star) is routinely inoculated with phytoplasma in commercial nurseries resulting in highly branched plants desirable for sale (Lee *et al*. 1997). However, phenotype variation (Lee *et al*. 2021), loss of phytoplasma at higher temperatures, i.e., > 35°C (Linck *et al*. 2019), and tedious manual infection of phytoplasma with increased transmission risk of pathogens such as viruses call for more sustainable approaches including breeding and genetic engineering towards higher branching of non-infected plants. Poinsettia is amenable for genetic manipulation, for example, poinsettia flavonoid 3-hydroxylase (Vilperte *et al*. 2019) has been knocked out for bract color change from red to orange (Nitarska *et al*. 2021). Here, we aimed to identify candidate regulatory genes that could serve as targets for future genetic engineering efforts toward phytoplasma-free, branching poinsettia plants. In this study, transcriptome assembly and expression analyses in phytoplasma-infected and non-infected poinsettia plants identified two genes, coding for MAF3 (a MADS-box protein) and bZIP67 transcription factors that may play roles in branching. Our results were further confirmed by demonstrating that *Arabidopsis maf*3 and *bzip*67 mutants have highly branching phenotypes. Finally, genome-wide transcriptome analysis of these *Arabidopsis* mutants revealed possible down-stream targets and transcriptional regulatory effects. The two transcription factors could thus be promising candidates for development of stable phytoplasma-free high branching lines of poinsettia.

## Materials and methods

### Plants

Poinsettia (*E. pulcherrima*) cv. EXP 98 plants were obtained from Graff Breeding A/S, Sabro, Denmark. At Graff Breeding, a phytoplasma-free (EXP 98-1) plant was infected with phytoplasma (*C.* Phytoplasma pruni) via grafting (EXP 98-2). Cuttings of both types (EXP 98-1 and EXP 98-2) were kept in a greenhouse at 16 hours light cycles and 22-24°C and trimmed two months before sample collection. Buds number two, three and four from the top of the stems were collected from phytoplasma-infected (EXP98-2) and non-infected (EXP 98-1) plants. In total, 30 buds were isolated from each type (three replicates each with pooled samples from 10 plants). These buds were selected as they were at the initial phase of growth.

*Arabidopsis* seeds for wild-type and mutant lines were obtained from the European Arabidopsis Stock Center (NASC at https://arabidopsis.info/BasicForm; *Arabidopsis maf*3 NASC ID: N674795, *Arabidopsis bzip*67 NASC ID: N666597, *Arabidopsis brc*1/*tcp*18 NASC ID: N857231, wild-type *Arabidopsis* Col-2 NASC ID: N907, and wild-type *Arabidopsis* Col-0 NASC ID: N70000). Mutant lines were described previously (Alonso *et al*. 2003; Woody *et al*. 2007). The *Arabidopsis brc*1/*tcp*18 is obtained from Col-2 ecotype and, *bzip*67 and *maf*3 mutant lines are obtained from Col-0 ecotype. For vernalization, seeds were distributed in petri-dishes covered with moist paper and kept in darkness at 4-5°C for eight days. The seeds were planted in pre-watered pots and kept in a climate chamber (21°C, 60% humidity, 16 hours light [(100 µmol(µE) m^-2^ s^-1^] and eight hours night (20°C, 60% humidity) until germination. For promoting vegetative growth, germinated plants were kept in short-day cycles (21°C, 50% humidity, 10 hours light [(70 µmol(µE) m^-2^ s^-1^)] and 14 hours night (21°C, 50% humidity) for three weeks. Finally, the plants were transferred to long-day cycles of 16 hours light (100 µmol(µE) m^-2^ s^-1^, 22°C, 50% humidity) and eight hours night (20°C, 50% humidity). All plants, one plant per pot, were bottom-watered. We were interested in the number of primary branches as it is important for poinsettia bushy phenotype; cauline branches are dependent on the stem length and show high within-line variability. We therefore examined the number of primary branches in a minimum of 10 plants from each *Arabidopsis* line to compare the wild-type and mutant lines. This was performed two and four weeks after inflorescence emergence (*Arabidopsis* growth stage 5.0, see (Boyes *et al*. 2001); ≈ the end of week 6 and 8 after germination). We included all the emerged primary branches, i.e., of any length, for each plant, grown individually in pots. Col-0 and Col-2 wild-type lines were included in the experiment for comparison.

### Sample collection for genomic DNA and total RNA extraction

Genomic DNA from poinsettia plants for phytoplasma quantification were extracted from petioles and leaf-midribs. Genomic DNA for *Arabidopsis* plants were obtained from leaves. For RNA extraction from poinsettia plants, bud number two, three, and four from the top of stems were collected. Due to the small sizes, bud number two and three (bud#2/3) were pooled before RNA extraction to have enough material. *Arabidopsis* RNA samples were extracted from leaves and siliques. The reason to focus on the leaves and siliques and not on the buds was the impracticality of sampling enough and synchronized tissue from very tiny-size buds of *Arabidopsis*. Challenges in sample collection can markedly affect reproducibility in transcriptome analysis, even with the use of advanced techniques such as laser-capture microdissection. Leaves and siliques were chosen because *Maf*3 and *bZip*67 have the highest expressions in these tissues (see the Result section). Sample collection was performed when *Arabidopsis* plants had siliques at stage 4-6 as defined by Mizzotti et al. (Mizzotti *et al*. 2018). Transcriptome analyses were performed to investigate the possible transcriptional regulatory effects of the *maf*3 *and bzip*67 mutants rather than for direct study of the possible branching-specific regulatory effects. All samples were immediately flash frozen in liquid nitrogen and kept at -80°C until processed. We used sterile mortars and pestles for tissue grinding in liquid nitrogen to extract genomic DNA using the DNeasy Plant Mini kit (QIAGEN). DNA concentrations were quantified by a Qubit 3 fluorometer (Invitrogen). For RNA extraction, buds were first washed with 70% cold (-80°C) acetone to remove milky sap. RNeasy Plant Mini kit (QIAGEN) was used for RNA extraction from poinsettia and *Arabidopsis* plants. On-column DNA digestion was carried out for 20 minutes at room temperature using the RNase-Free DNase (Cat. No. 79254; Qiagen). Quantity and quality of RNA samples were examined by an Agilent 2100 Bioanalyzer and its associated RNA 6000 Nano assay kit (Cat. No. 1222; Agilent Technologies, see Fig. S1).

### RNA sequencing and quality check

High-quality RNA samples were used for TruSeq stranded mRNA paired end (2×101) sequencing by NovaSeq 6000 by Macrogen, Netherlands. Reads were quality checked using FastQC (Version 0.12.0). Based on the overall quality, we trimmed the reads using Trimmomatic (version 0.39) (Bolger *et al*. 2014). First, 12 nucleotides from 5-prime ends were removed. This was followed by trimming nucleotides with quality scores below five (i.e., error likelihood less than 0.31623) from both ends and trimming four-nucleotide sliding windows if the average quality was below six (i.e., error likelihood less than 0.25119). Only surviving paired reads with minimum 46 nucleotides in length (2 × ≥ 46) were kept for downstream analyses.

### De novo assembly of the poinsettia transcriptome and remapping reads for expression analysis

The read headers were adjusted within the fastq files and using cut and perl commands to have pairs recognizable by the Trinity tool (Grabherr *et al*. 2011; Haas *et al*. 2013). For the poinsettia transcriptomes, we performed strand specific de novo assembly (with options –min_kmer_cov 2 & no-normalize_reads) using Trinity (Grabherr *et al*. 2011; Haas *et al*. 2013), Bowtie2 (Langmead & Salzberg 2012 2; Langmead *et al*. 2019) and samtools (Li *et al*. 2009; Danecek *et al*. 2021). Benchmarking Universal Single-Copy Orthologs (BUSCO) (Manni *et al*. 2021; Tegenfeldt *et al*. 2025) were applied to assess the completeness of the de novo assembled poinsettia transcriptome. BUSCO was performed using different lineages including eukaryotes, eudicots (all dicot plants), the brassicales order, and the fabales order. The taxonomic tree of the lineages was obtained by the Taxallnomy web application (Sakamoto & Ortega 2021). *For Arabidopsis thaliana* transcriptome analysis, we used Bowtie2 (Langmead & Salzberg 2012; Langmead *et al*. 2019), samtools (Li *et al*. 2009; Danecek *et al*. 2021), and RSEM (Li & Dewey 2011) to build the index reference transcriptome based on the *Arabidopsis* TAIR11 assembly. These transcripts cover 5UTR, exons, and 3UTR. Afterwards, we added a poly A sequence of 150 bases to the end of every transcript. The QC passed reads were mapped, by Bowtie2 (Langmead & Salzberg 2012; Langmead *et al*. 2019), onto 58,203 processed and indexed transcripts representing 38,086 genes.

### Gene expression analysis

After remapping the reads to the assembled poinsettia transcripts, RSEM (Li & Dewey 2011) was used to estimate gene expression levels. The bias levels were examined as previously introduced (Darbani *et al*. 2014, 2015) to shed light on the inter-treatment bias for cell-numbers. The analysis was based on 17 reference genes from poinsettia selected as the orthologues of 13 most stable reference genes from rice (Darbani *et al*. 2014). The reference genes were tubulin alpha-1 chain (TRINITY_DN42_c0_g1, TRINITY_DN22410_c2_g1), actin 1 (TRINITY_DN5365_c0_g1, TRINITY_DN212_c0_g1), ubiquitin (TRINITY_DN630_c0_g1), eukaryotic initiation factor 5c (TRINITY_DN847_c0_g1), Splicin factor U2af (TRINITY_DN855_c0_g1, TRINITY_DN6067_c0_g1), formin-binding protein (TRINITY_DN7868_c0_g1), protein kinase (TRINITY_DN10678_c1_g1), polyubiquitin (TRINITY_DN3355_c0_g1, TRINITY_DN30_c6_g1, TRINITY_DN999_c0_g2, TRINITY_DN30_c0_g1), translationally-controlled tumor protein (TRINITY_DN3896_c0_g1), SKP1-like protein 1A (TRINITY_DN545_c0_g3), and GADPH (TRINITY_DN86_c0_g1). The raw expression count data was used for differential analysis using edgeR (Robinson *et al*. 2010; Chen *et al*. 2025 4). The fold-change threshold levels were determined to have less than 5% or 1% likelihood to be surpassed by inter-replicate fold-changes (random fold-changes) as previously reported (Darbani & Stewart 2014). The same analysis was performed for *Arabidopsis* data using *Arabidopsis* stable reference genes that were reported previously (Czechowski *et al*. 2005; Dekkers *et al*. 2012; Kudo *et al*. 2016; Ferreira *et al*. 2023). This included AT1G07920, AT1G07930, AT1G11650, AT1G13320, AT1G13440, AT1G16970, AT1G50010, AT1G59830, AT2G28390, AT2G43770, AT3G12210, AT3G18780, AT3G26520, AT3G42050, AT4G05320, AT4G10790, AT4G12590, AT4G27960, AT4G40030, AT5G15710, AT5G25760, AT5G42000, AT5G53300, AT5G55840, AT5G56290, and AT5G60390. To obtain gene expression heatmaps across samples, the expression levels were used to measure the Pearson correlation matrix for pairwise sample comparisons based on the selected set of differentially expressed features (Grabherr *et al*. 2011; Haas *et al*. 2013). Metascape (Zhou *et al*. 2019) was used for pathway enrichment analysis of the differentially expressed genes. We applied the ForceAtlas2 graph algorithm for protein-protein network interaction analysis (Jacomy *et al*. 2014) with the gravity set to 0.8. The experimentally confirmed protein-protein and protein-DNA interactions were extracted from *Arabidopsis* Interactions Viewer 2 (AIV2) (Geisler-Lee *et al*. 2007; Dong *et al*. 2019).

### Annotation of the differentially expressed genes

ORFs were mapped using TransDecoder-v5.7.0 (Haas *et al*. 2013) and a minimum peptide length of 50 amino acids. The longest ORFs of every gene were kept for downstream analyses. Protein motif identification was carried out using HMMER (v3.3.2) (Eddy 2011) and based on Pfam34.0 motifs (Mistry *et al*. 2021). The UniProt proteome data (Bryant *et al*. 2017; The UniProt Consortium 2025) as well as *Arabidopsis* proteome (TAIR11) were indexed and used as reference databases to BLAST (blastx; evalue < 1e-5) for the homologues of the assembled poinsettia transcripts. We used the NCBI Blast machine executables version 2.13.0 (available at https://blast.ncbi.nlm.nih.gov/doc/blast-help/downloadblastdata.html). Trinotate-v3.2.2 (Bryant *et al*. 2017) was finally applied to merged annotations.

### Phylogenetic analysis

*Arabidopsis* bZIP transcription factor family members, poinsettia bZIP67, and the AtbZIP67/EpbZIP67 homologues from *Lotus japonicus*, *Hibiscus syriacus*, *Prunus persica*, *Populus euphratica*, *Gossypium hirsutum*, *Theobroma cacao*, *Manihot esculenta*, *Jatropha curcas*, *Mercurialis annua*, *Ricinus communis*, and *Brassica napus* were examined for evolutionary trends. We also performed phylogenetic analysis on the MAF1‒5 and other AGAMOUS protein family members including AG, AGL20, AGL10, SEPALLATA2, APETALA1, FLC, FYF, and TT16 from *Arabidopsis* and their homologues from poinsettia, *Solanum lycopersicum*, *Vitis vinifera*, *Nicotiana tabacum*, *Populus trichocarpa*, *Ricinus communis*, *Manihot esculenta*, *Camelina sativa*, *Capsella rubella*, *Brassica napus*, and *Theobroma cacao*. Alignments were executed with Gap Opening Penalty of 10, Extension Penalty of 0.2, 30% Delay Divergent Cutoff, and a positive-value adjusted distance matrix using the tool MEGA11 and its alignment module ClustalW (Tamura *et al*. 2021). Phylogenetic model selection and tree construction were carried out using the iqtree v1.6.12 (Nguyen *et al*. 2015). Tree visualization was performed by FigTree v1.4.4 (available at http://tree.bio.ed.ac.uk/software/figtree/).

Bootstrapping was set to 1000 and performed through two approaches including the approximate likelihood-ratio test (aLRT; based on the log ratio between the likelihood value of the current tree and that of the best alternative) (Guindon *et al*. 2010) and the ultrafast bootstrap algorithms (Minh *et al*. 2013).

### Screening of the plants

To screen for phytoplasma infection, a Taq-Man assay was performed by adding primers, probe, and template to the master mix of qPCRBIO Probe Blue Mix Lo-ROX (PCR Biosystems Ltd.) as previously described (Christensen *et al*. 2004). We used an AB applied biosystems ViiA7 thermocycler. The primers and amplification condition were as previously reported (Christensen *et al*. 2004). The *Arabidopsis* lines were PCR-screened for the existence of T-DNA insertion at the expected genomic positions. PCR reactions were performed using 10 ng template genomic DNA, 3% DMSO, and the Phusion U Hot Start DNA Polymerase (Thermo Scientific). Initial denaturation was at 95°C for four min followed by 98°C for 30 sec. Afterwards, 40 cycles of amplification (98°C for 10 sec, 71°C for 40 sec, 72°C for 40 sec) were executed. For end-repair, a final extension step of 10 min at 72°C was also included. The primer sequences for PCR screening were as followed: for *maf*3 mutant (MAF.r2: 5ʹAAGTACTTGAACAGCATTGAGAATGTATCAAC3ʹ, MyLB: 5ʹCGATTTCGGAACCACCATCAAACAGGAT3ʹ), for *bzip*67 mutant (bZIP.r2: 5ʹCTTTCTCAATTTGTCTCCGCTCTTCTCTT3ʹ, MyLB: 5ʹCGATTTCGGAACCACCATCAAACAGGAT3ʹ), and for *brc*1 mutant (BRC.f2: 5ʹAGTACGTCGCCTGTGTTGTCTTTCTCA3ʹ, MyWiscLB2: 5ʹAGCGTCAATTTGTTTACACCACAATATATCCT3ʹ)

## Results

### Transcriptome assembly for poinsettia

We first aimed to assemble the transcriptome of axillary buds in poinsettia due to the lack of an annotated reference genome. Phytoplasma-free and -infected poinsettia plants (Fig. 1a) were used for axillary bud sampling and RNA extraction. Although the architecture of the plants was indicative of phytoplasma-free and -infected plants, this was confirmed by a TaqMan assay targeting the phytoplasma 16S gene (Fig. 1b). RNA extraction for transcriptome analysis was carried out for bud number two and three (pooled) and bud number four from the top of the stems (Fig. S2). First, we pooled the sequencing results from all samples to assemble a bud specific poinsettia transcriptome. More than 546 million paired-reads (Table S1) were assembled. We built 127,338 genes in the form of 213,118 transcripts (Table S2). The median and average transcript length were 502 and 847.4 nucleotides, respectively (Table S2). The N50 for the assembly was 1,398 nucleotides including up to 39,381 transcripts (Table S2).

**Fig. 1.**
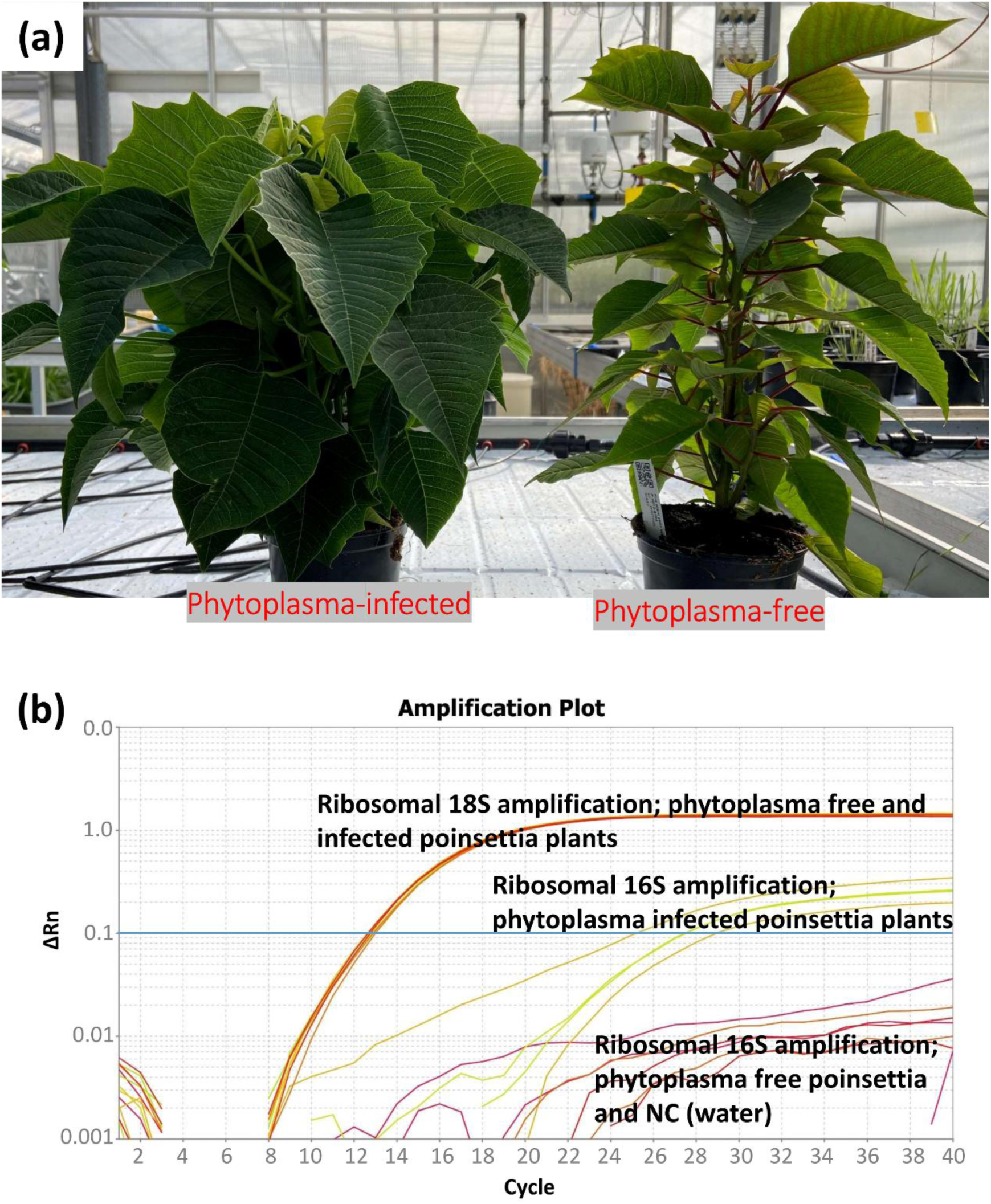
Poinsettia plants were examined for infection by phytoplasma. (a) Phytoplasma-free and -infected poinsettia plants. (b) TaqMan assay performed on plant genomic DNA for amplification of plant ribosomal 18S and phytoplasma 16S DNA. Assays were performed in duplicates on two phytoplasma-infected and two phytoplasma-free poinsettia plants. ΔRn: The magnitude of the signal generated, Rn: Normalized reporter or passive reference dye-based corrected fluorescence emission intensity.

These results, i.e., average and median length as well as N50, were indicative of a high-quality transcriptome assembly. In our bud-specific transcriptome assembly, there were 39,381 transcripts with a minimum length of 1,398 nucleotides. The latest *Arabidopsis* assembly (TAIR11; https://www.arabidopsis.org/) codes for 48,266 transcripts (average ORF+3ʹUTR length = 1,574 nucleotides) of which 23,478 transcripts have a minimum ORF+3ʹUTR length of 1398 nucleotides. Using long-read Nanopore RNA-sequencing, two previous studies in *Arabidopsis* have also reported a read N50 of ≈1200 nucleotides (Zhang *et al*. 2020; Wang *et al*. 2024). Furthermore, the assembled poinsettia transcriptome covered all eukaryotic core genes as examined by BUSCO (Fig. 2b, see Materials and methods for details) which is an indication of a full-length transcriptome assembly. Poinsettia is a dicot plant in Euphorbiaceae which has Brassicaceae and Fabaceae as the closest families with available genetic data (Fig. 2a) and, as expected, we found 2.5%, 17.9%, and 22% missing transcripts within the assembly by narrowing down through the phylogenetic tree and examining ‘dicots’, ‘Brassicales’, and ‘Fabales’, respectively (Fig. 2b). This increase in missing transcripts is due to the inclusion of less phylogenetically conserved genes. In addition, BLAST analysis performed against the *Arabidopsis* proteome revealed that the assembled poinsettia bud-specific transcriptome represents up to 15,214 *Arabidopsis thaliana* proteins with higher than 50% coverage (Fig. 2c). Further analysis revealed that 98% of the total gene expression belonged to the top 65,268 highly expressed genes while the 62,070 lowest expressed genes only contributed to the 2% of the total expression level (Fig. 2d). Overall, our results indicated a high quality transcriptome assembly in terms of transcript completeness and overlap rate with the *Arabidopsis* transcriptome as compared to the previously reported bract transcriptome of the poinsettia (Vilperte *et al*. 2019).

**Fig. 2.**
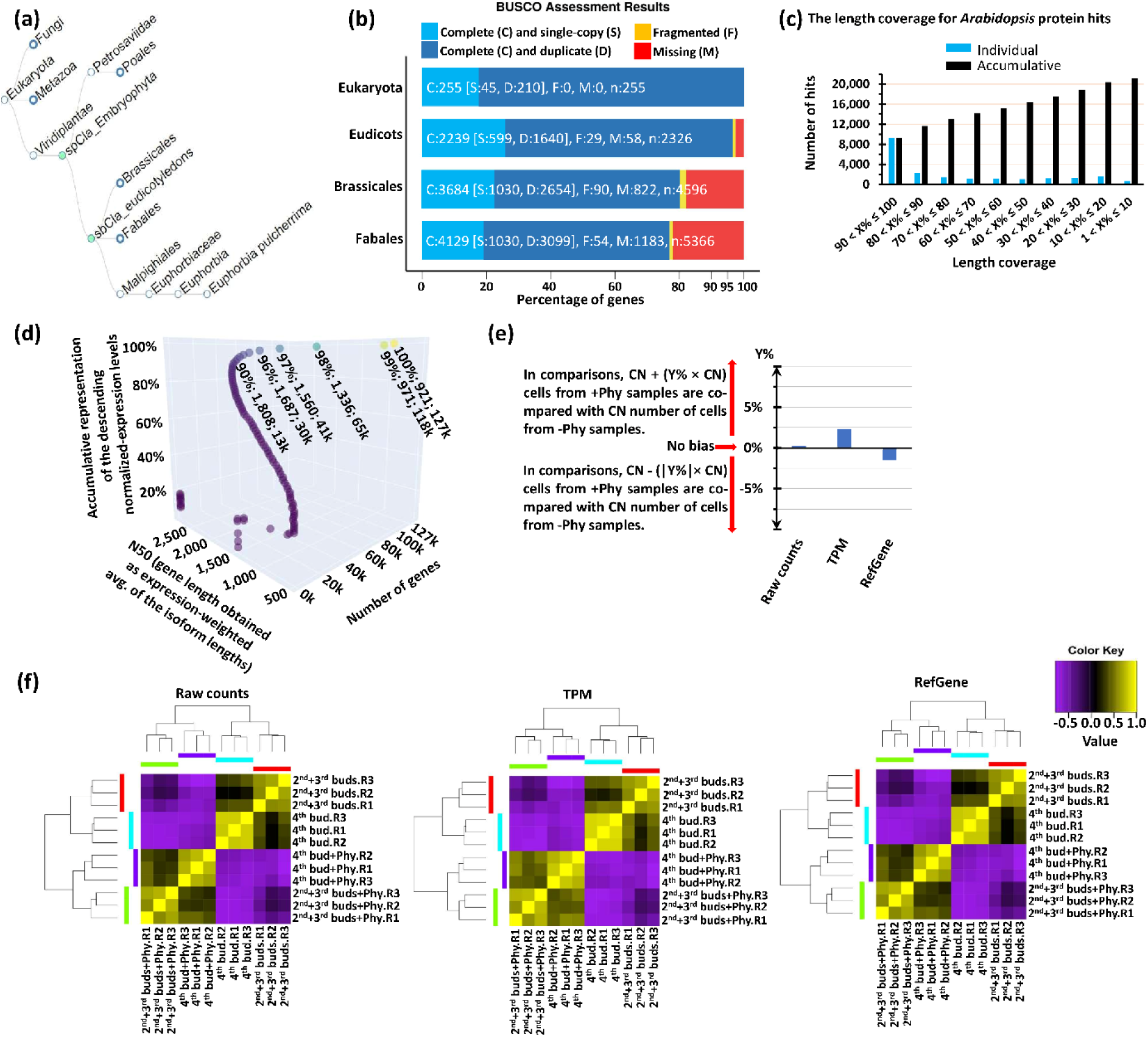
Assembly characteristics and expression data normalization. (a-d) The completeness of the assembled poinsettia transcriptome. (a) A taxonomic tree is drawn by the Taxallnomy web application (Sakamoto and Ortega, 2021). (b) Percentage of examined core genes with or without orthologue genes in poinsettia. Orthologues are categorized as complete/partially assembled or as single/multi-copy genes. (c) The assembled poinsettia transcriptome represents more than 15,240 *Arabidopsis* proteins found with a length coverage above 50% after Blast analysis against the *Arabidopsis thaliana* proteome. (d) Up to 65,268 (51.26%) genes were represented by 98% of the assembled sequencing reads and the left 2% were assembled into 62,070 (48.74%) weakly expressed genes. (e,f) Inter-treatment bias levels for cell-number. (e) The bias for cell-number was examined while comparing the phytoplasma-free with the phytoplasma-infected plants (+Phy/-Phy). (f) Heatmap of the sample clusters based on the expression levels of 3900 differentially expressed (DE) genes (FDR < 0.001, FC ≥ 2.5). First, DE genes were identified on raw count data using edgeR and afterwards, raw count data, TPM (transcript per million reads) data, and reference gene (GADPH coding gene) based corrected data were used to obtain the Pearson correlation heatmaps. CN represents the quantity for number of cells. R1/R2/R3: replicates.

### Quantification of expression in phytoplasma-free and -infected poinsettia buds

While genome-wide normalization methods assume no difference in total cellular RNA content among the samples, we could, however, expect different levels of total cellular RNA content among tissues and treatments. Genome-wide normalization approaches cannot deal with this difference in total cellular RNA content and can lead to major biases. Therefore, we examined the bias for cell number as described previously (Darbani *et al*. 2014), but found no considerable bias, i.e., < 3% (Fig. 2e). This agreed with the heatmap of the sample clusters which were obtained based on the expression level correlations of differentially expressed genes and showed full alignment among the raw and corrected expression data (Fig. 2e,f). The raw expression count data was therefore used for differential analysis as there was no considerable bias among the samples as well as among the normalization approaches (Fig. 2e,f).

By performing differential analysis for the combined buds two and three (bud#2/3), we found 236 and 298 genes with known homologues that were up- or downregulated in phytoplasma-infected tissues, respectively (FDR < 0.001; Fig. 3a, Table 1). After applying an inter-replicate based fold-change threshold of less than 5% and 1% likelihood to be surpassed by inter-replicate fold-changes (random fold-changes) as explained previously (Darbani & Stewart 2014), we retained 169 (2.5 ≤ FC < 5.2; FDR < 0.001) and 18 (5.2 ≤ FC; FDR < 0.001) up- or downregulated genes, respectively (Fig. 3a). When analyzing bud number four (bud#4) in phytoplasma-free and -infected plants, we found 1,685 and 1,902 genes (FDR < 0.001) with known homologues that were upregulated or down-regulated, respectively (Fig. 3, Table 1). After applying an inter-replicate-based fold-change threshold with less than 5% and 1% likelihood to be surpassed by inter-replicate fold-changes (random fold-changes), we found 1224 (2.9 ≤ FC < 9.1; FDR < 0.001) and 95 (9.1 ≤ FC; FDR < 0.001) differentially expressed genes, respectively (Fig. 3a). The differentially expressed genes with no homologues were notably predicted to code for peptides shorter than 100 amino acids (Fig. S3). Pathway enrichment analysis on all the known, i.e., with homologues, differentially expressed genes revealed cell cycle and tissue developmental modulations as well as phenylpropanoid, flavonoid, and brassinosteroid biosynthesis pathways (Fig. 3b). Taken together, there was a good agreement between bud#2/3 and bud#4 as the majority of the differentially expressed genes in bud#2/3 were also found differentially expressed in bud#4 (Table 1).

**Fig. 3.**
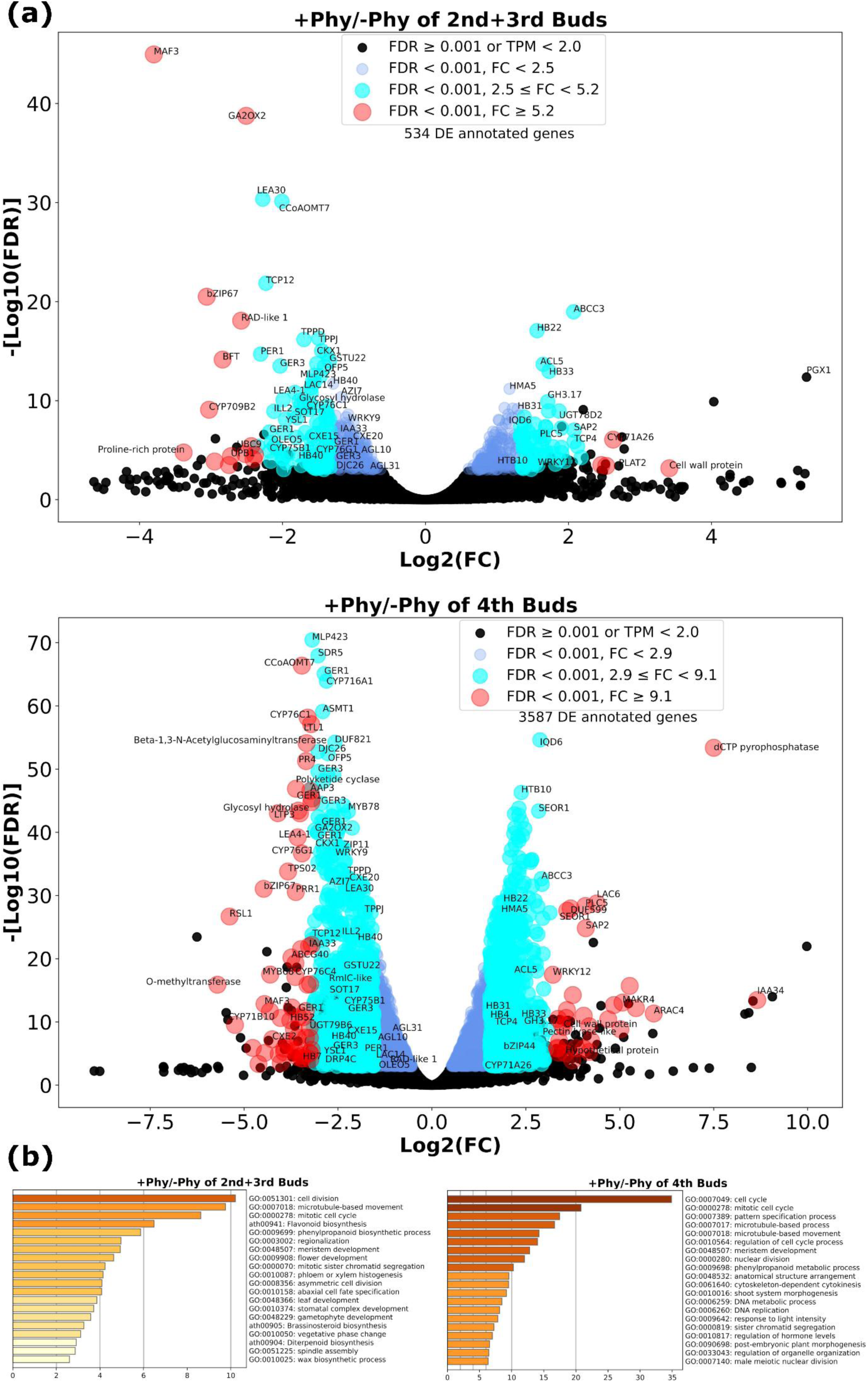
Differentially expressed poinsettia genes. (a) Volcano plots of gene expression levels in phytoplasma-free and - infected poinsettia plants. Fold-change (FC) thresholds were determined to have less than 5% (FC for bud#2/3 ≥ 2.5, FC for bud#4 ≥ 2.9) or 1% (FC for bud#2/3 ≥ 5.2, FC for bud#4 ≥ 9.1) likelihood to be surpassed by inter-replicate fold-changes, i.e., random fold-changes, as reported previously (Darbani & Stewart, 2014). (b) Pathway enrichment on the *Arabidopsis* homologues of the differentially expressed genes, i.e., FDR < 0.001 and TPM ≥ 2.

**Table 1.** Number of differentially expressed genes between phytoplasma-infected and -free poinsettia plants.

|  | Samples |  | Shared among samples |
| --- | --- | --- | --- |
|  | bud#2/3 | bud#4 |  |
| Annotated differentially expressed genes (+Phy/-Phy) |  |  |  |
| Upregulated | 236 | 1,685 | 213 |
| Downregulated | 298 | 1,902 | 255 |
| Sum | 534 | 3,587 | 468 |
| Unknown differentially expressed genes (+Phy/-Phy) |  |  |  |
| Upregulated | 62 | 290 | 50 |
| Downregulated | 135 | 355 | 110 |
| Sum | 197 | 645 | 160 |

The observed considerably small biases, i.e., ≤ 5%, and also use of reference gene based normalization eliminated the need for validation of the expression quantities through real-time PCR. Real-time PCR has the advantage of comparing equal number of cells between the samples due to the use of reference gene based normalization. But it also has the disadvantages of indirect measurements, non-specific amplifications, and being dependent on the amplification rates. In contrast, RNA-Sequencing benefits from direct measurements (counting of the sequencing reads) and when coupled with reference gene based normalization, can also ensure the equality of number of cells being compared among the samples (Darbani *et al*. 2014). Therefore, RNA sequencing using sufficient starting RNA material, which eliminates the need for an initial amplification step, provides higher accuracy than qPCR. Furthermore, validating RNA-Sequencing results by real-time PCR is only reasonable and necessary when the reference gene based correction is not a part of the normalization of raw quantities showing considerable level of bias for cell number, e.g., ≥ 25%.

### Top differentially expressed genes in poinsettia buds in response to phytoplasma infection

Since our interest lies in bud dormancy rather than stem development and elongation, we focused solely on comparing phytoplasma-free and phytoplasma-infected poinsettia plants, without comparing bud #2/3 and bud #4. The expression levels of the genes and fold changes between the phytoplasma-free and -infected poinsettia plants are available in Dataset S1. To facilitate future genetic engineering efforts in poinsettia, our focus was on identifying candidate genes encoding regulatory factors that are repressed during phytoplasma infection. These factors can produce stable phenotypes through their wide down-stream regulatory effects, and their transcription suppression is an easy target through CRISPR-Cas gene editing technology.

When compared to the phytoplasma-free plants, the gene with the highest repression in the phytoplasma-infected plants (14- and 22-fold in bud#2/3 and bud#4, respectively) was *EpMaf*3, a homologue of the *Arabidopsis Maf*3 gene (AT5G65060; Fig. 3a) that has been shown to be under positive selection and diversification (Caicedo *et al*. 2009). AtMAF3 is also involved in flowering repression in coordination with other flowering repressors such as Flowering locus C (Gu *et al*. 2013). As illustrated in Fig. 3a, we also noticed *Ep*b*Zip*67/*Dpbf*2 (with *Arabidopsis* homologue AT3G44460) among the top differentially repressed genes. The *Ep*b*Zip*67 gene was repressed eight-fold in bud#2/3 and 22-fold in bud#4 in response to phytoplasma infection. The functional roles of bZIP transcription factors include the regulation of different processes such as defense against pathogens, abiotic stress, and developmental signaling (Dröge-Laser *et al*. 2018). *Arabidopsis* bZIP67 has previously been found to be involved in fatty acid metabolism in seeds (Mendes *et al*. 2013 3), and in seed dormancy (Bryant *et al*. 2019). A third interesting candidate was *EpCyp*709B2 (with *Arabidopsis* homologue AT2G46950) which showed eight- and 15-fold repression in bud#2/3 and bud#4 of phytoplasma-infected plants, respectively (Fig. 3a). We further noticed two *EpHb*40 genes (with *Arabidopsis* homologue AT4G36740) which were down-regulated two- to four-fold in bud#2/3 and four to six fold in bud#4 in phytoplasma-infected plants when compared to the phytoplasma-free plants. The HOMEOBOX genes *EpHb*7 (with *Arabidopsis* homologue AT2G46680) and *EpHb*52 (with *Arabidopsis* homologue AT5G53980) were down-regulated 10- and 14-fold, respectively, by phytoplasma infection in bud#4 (Fig. 3a). In contrast, we found other HOMEOBOX coding genes such as *EpHb*4, 22, 31, and 33 that were up-regulated by phytoplasma infection (Fig. 3a). In agreement with the down-regulation of two *EpHb*40 genes, the suppression of branching via abscisic acid accumulation is facilitated by *Brc*1 and *Hb*40 genes (González-Grandío *et al*. 2017). BRC1/TCP18 is a well-known repressor of branching (Aguilar-Martínez *et al*. 2007) and the phytoplasma SAP11 effector protein destabilizes TCP transcription factors such as BRC1 through direct interaction to release branching (Sugio *et al*. 2011; Chang *et al*. 2018; Pecher *et al*. 2019). Here, we also report a down-regulation at transcript level of *Ep*BRC2/TCP12, triggered by phytoplasma infection. We found the gene coding for *Ep*BRC2/TCP12 (with *Arabidopsis* homologue AT1G68800 which is the closest homologue of BRC1 (Martín-Trillo & Cubas 2010)) with five-fold repression in bud#2/3, and nine-fold repression in bud#4 of the phytoplasma-infected plants when compared to the phytoplasma-free poinsettia plants (Fig. 3a). In contrast, the *EpTcp*4 gene was up-regulated four-fold in bud#2/3 and, three-fold in bud#4 in response to phytoplasma infection (Fig. 3a). In agreement, *EpTcp*4 has been found to be involved in shoot regeneration (Yang *et al*. 2020). Two gibberellin 2-beta-dioxygenase 2 (*EpGa2ox*2) genes (with *Arabidopsis* homologues AT1G30040 and AT1G02400), involved in active-to-inactive gibberellin conversion and thereby in inhibition of branching (Ni *et al*. 2015; Katyayini *et al*. 2020), were also repressed up to six-fold in bud#2/3 and nine-fold in bud#4 of phytoplasma-infected plants when compared to the phytoplasma-free plants (Fig. 3a). The regulatory role of sugar deprivation on axillary bud dormancy is discussed in the introduction. We accordingly found that the trehalose 6-phosphate phosphatase J gene (*EpTppj*; *Arabidopsis* homologue AT5G65140) was repressed three-fold in bud#2/3 and 3.5-fold in bud#4 of phytoplasma-infected plants in comparison with phytoplasma-free plants (Fig. 3a). The trehalose 6-phosphate phosphatase D gene (*EpTppd*; with *Arabidopsis* homologue AT1G35910) was also repressed three- and five-fold by phytoplasma infection in bud#2/3 and in bud#4, respectively (Fig. 3a). Although the *Arabidopsis* carboxylesterase 15 and 20 (*Cxe15 and* 20) enhance branching through strigolactone sequestration and hydrolysis (Roesler *et al*. 2021; Xu *et al*. 2021), we found three *EpCxe* genes (*Cxe*2, 15, and 20) repressed by phytoplasma infection (Fig. 3a). Finally, we found the auxin signaling negative regulators (Yu *et al*. 2022) of IAA33 (with *Arabidopsis* homologue AT5G57420) down-regulated by phytoplasma infection. In parallel, auxin biosynthesis (Zheng *et al*. 2016; Yu *et al*. 2022) was activated: two genes coding for indole-3-pyruvate monooxygenase YUCCA6 (with *Arabidopsis* homologue AT5G25620) and indole-3-acetic acid-amido synthetase gene GH3.17 (with *Arabidopsis* homologue AT1G28130) were upregulated in phytoplasma-infected plants (Fig. 3a). Although auxin signaling, initiated in the shoot tip, represses axillary branching, the induction of *Yucca*6 and *Gh*3.17 in phytoplasma-mediated activated axillary buds may play an important role in establishing nascent shoots.

### Characterization of branching phenotypes and transcriptomic alterations in Arabidopsis maf3 and bzip67 mutant lines

We used publicly available *Arabidopsis* mutant resources as a reliable and efficient approach to evaluate the selected candidate genes. Although this approach alone has limited direct applicability to poinsettia, concordance between the *Arabidopsis* mutant results and poinsettia transcriptome analyses strengthens confidence in these candidate genes for future genetic engineering efforts. We conducted *in vivo* functional analysis on the top two transcriptional hits using *Arabidopsis thaliana* mutant lines. We selected the top down-regulated transcription factor genes *Maf*3 and *bZip*67 for examination of their roles in stem branching in *Arabidopsis*. Bud #2/3 was developmentally representative of earlier stages of dormancy release compared with bud #4. Therefore, the selection of candidate genes was primarily based on observations from bud #2/3 and their confirmation in bud #4. *Epmaf*3 showed the lowest *p*-value and the highest repression rate in bud #2/3. The candidate gene *bZip*67 ranked second in repression rate and was also among the six genes with the smallest *p*-values in bud #2/3. Notably, both genes were also among the five most highly repressed genes in bud #4. We also included the *Tcp*18 transcription factor gene as its role in repression of stem branching is well documented. In contrast to the well-documented role of TCPs in shoot branching, it is unknown how MAF3 and bZIP67 are involved in shoot branching. As shown in Fig. S8, the phylogenetic tree of MAFs and AGAMOUS like proteins, all from MADS-box transcription family, indicates that MAF members and Flowering Locus C (FLC) have recently diversified evolutionarily, and within taxonomic families. This, independently, also applies to AGAMOUS-like 10 (AGL10) and AGAMOUS-LIKE 7 (APETALA1). AGAMOUS-LIKE 20 (AGL20), Forever Young Flower (FYF, also known as AGL42), AGAMOUS, Transparent Testa16 (TT16, also known as AGL32), and AGAMOUS-like 4 (SEPALLA2) are, in contrast, diversified from each other before the plant families radiated (Fig. S8). The within-plant family evolution of MAFs was not observed for bZIP transcription factors. The phylogenetic analysis of bZIP transcription factors revealed that bZIP67 evolved early and before radiation of plant species within distinct taxonomic families (Fig. S8). Previous studies point to the involvement of *Arabidopsis* bZIP67 in seed fatty acid metabolism (Mendes *et al*. 2013 3) and dormancy (Bryant *et al*. 2019).

The T-DNA insertion mutants of *Arabidopsis maf*3 (NASC ID: N674795), *bzip*67 (NASC ID: N666597), and *brc*1/*tcp*18 (NASC ID: N857231) were used in the experiments. The T-DNA insertion is in intron II of *bZip*67 and in exon I of *Brc*1 and *Maf*3 (Fig. S4a). T-DNA insertions were confirmed by PCR (Fig. S4b). Resembling phytoplasma infection, we found a statistically significant increase (> two-fold; two-tailed t-test *p*-value < 6.7 × 10^-13^) in the number of primary shoots in all the mutant lines when compared to the wild-type lines (Fig. 4). Due to the lack of correlation between the length of primary inflorescence and the number of primary branches, between the length of cauline branches and the number of primary branches, and, also between the number of siliques and the number of primary branches (Fig. 4), examining the number of primary branches of plants at the same time, i.e., X days after germination or inflorescence emergence, is more accurate than counting the number of primary branches at different times among plants. Accordingly, we examined the number of branches two and four weeks after inflorescence emergence (Fig 4b, 4c, and Fig. S5) as described in the Methods section.

**Fig. 4.**
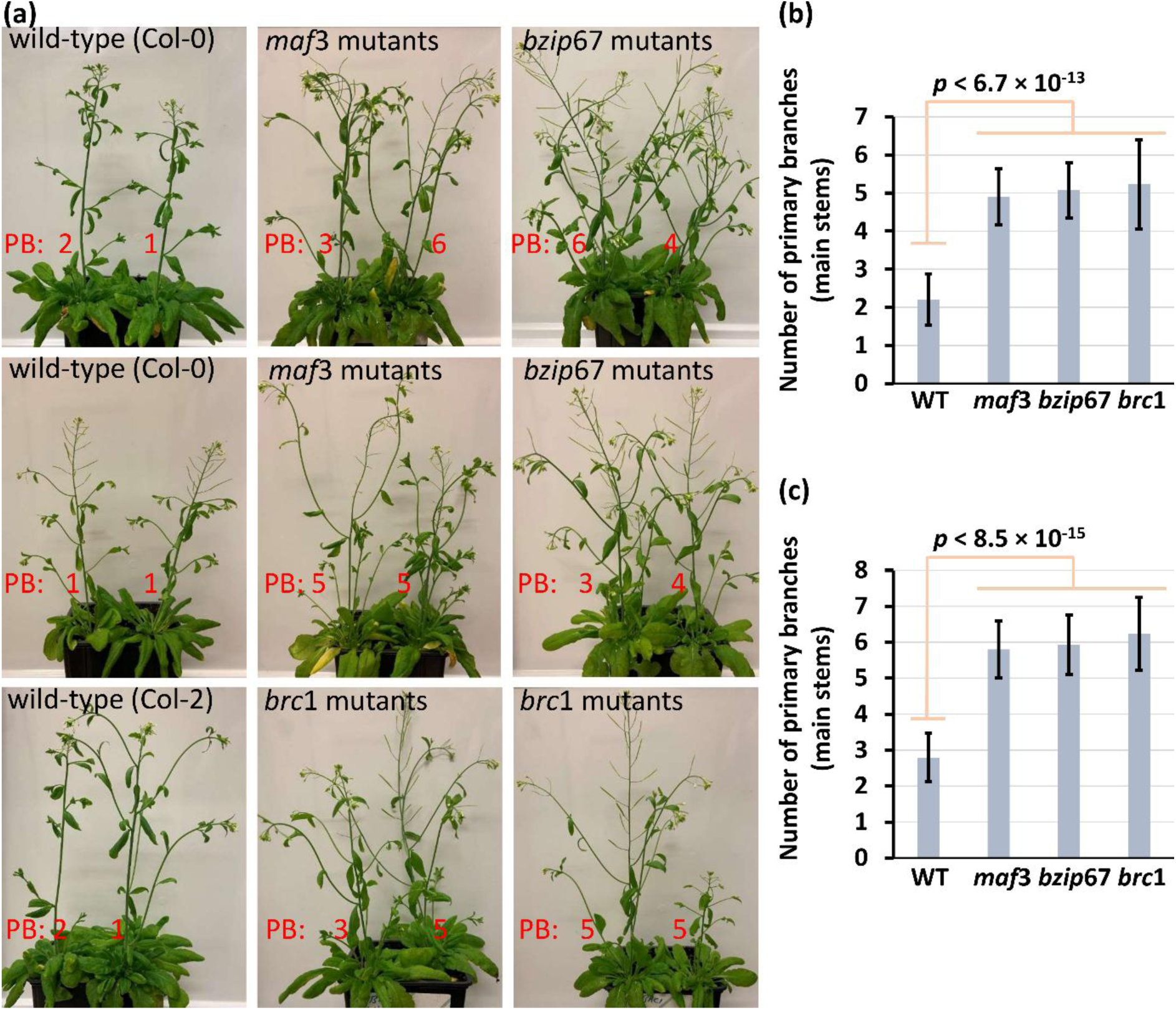
Number of primary branches (PB) in the wild-type and mutant *Arabidopsis thaliana* lines. (a) The number of primary branches (stems) are shown for two wild-type (Col-0 [NASC ID: N70000] and Col-2 [NASC ID: N907]) and three T-DNA insertion mutant lines (*maf*3 [NASC ID: N674795], *bzip*67 [NASC ID: N666597], and *brc*1 [NASC ID: N857231]). PB: number of primary branches. (b) The number of primary branches (stems) were examined two weeks after inflorescence emergence and in a minimum of 10 individual plants per line while growing one plant per pot (see Fig. S5). (c) The number of primary branches (stems) were examined four weeks after inflorescence emergence and in a minimum of 10 individual plants per line (one plant per pot). (b,c) Error bars represent standard deviations. All *p*-values from two-way t-tests were smaller than 6.7 × 10^-13^.

Next, to further investigate the possible transcriptional regulatory effects of the MAF3 and bZIP67 proteins rather than the possible branching-specific regulatory effects, we performed RNA-sequencing of the *Arabidopsis maf*3 and *bzip*67 mutants and their wild-types lines. Genome-wide transcriptome analyses were performed using RNA from the leaves of four *maf*3 plants and from the siliques of four *bzip*67 plants. By using the ePlant platform (Waese *et al*. 2017), we found that *AtbZip*67 and *AtMaf*3 are constitutively expressed. *AtMaf*3 shows relatively high expression across multiple tissues, whereas *AtbZip*67 is expressed at lower levels, with peak expression in young siliques. Therefore, we decided to use young siliques for examination of transcriptional perturbations in the *bzip*67 mutants.

As was the case in the poinsettia transcriptome analysis, the *Arabidopsis* samples showed less than 5% bias in cell-number (Fig. S6) and thereby, raw expression values were applied for differential expression analysis. The reported levels of bias for cell number are usually at higher rates, i.e., > 50% (Darbani *et al*. 2014). We found a few genes differentially expressed in the *maf*3 mutants when compared to wild-type *Arabidopsis* (Fig. 5, Dataset S1). Of most interest, *AtTcp*1 showed the highest up-regulation (78-fold) in the leaves of the *maf*3 mutants (Fig. 5, Dataset S1). Like *At*TCP18 (BRC1) and *At*TCP12 (BRC2), *At*TCP1 is also a member of the CYC-clade of TCPs (Martín-Trillo & Cubas 2010).

**Fig. 5.**
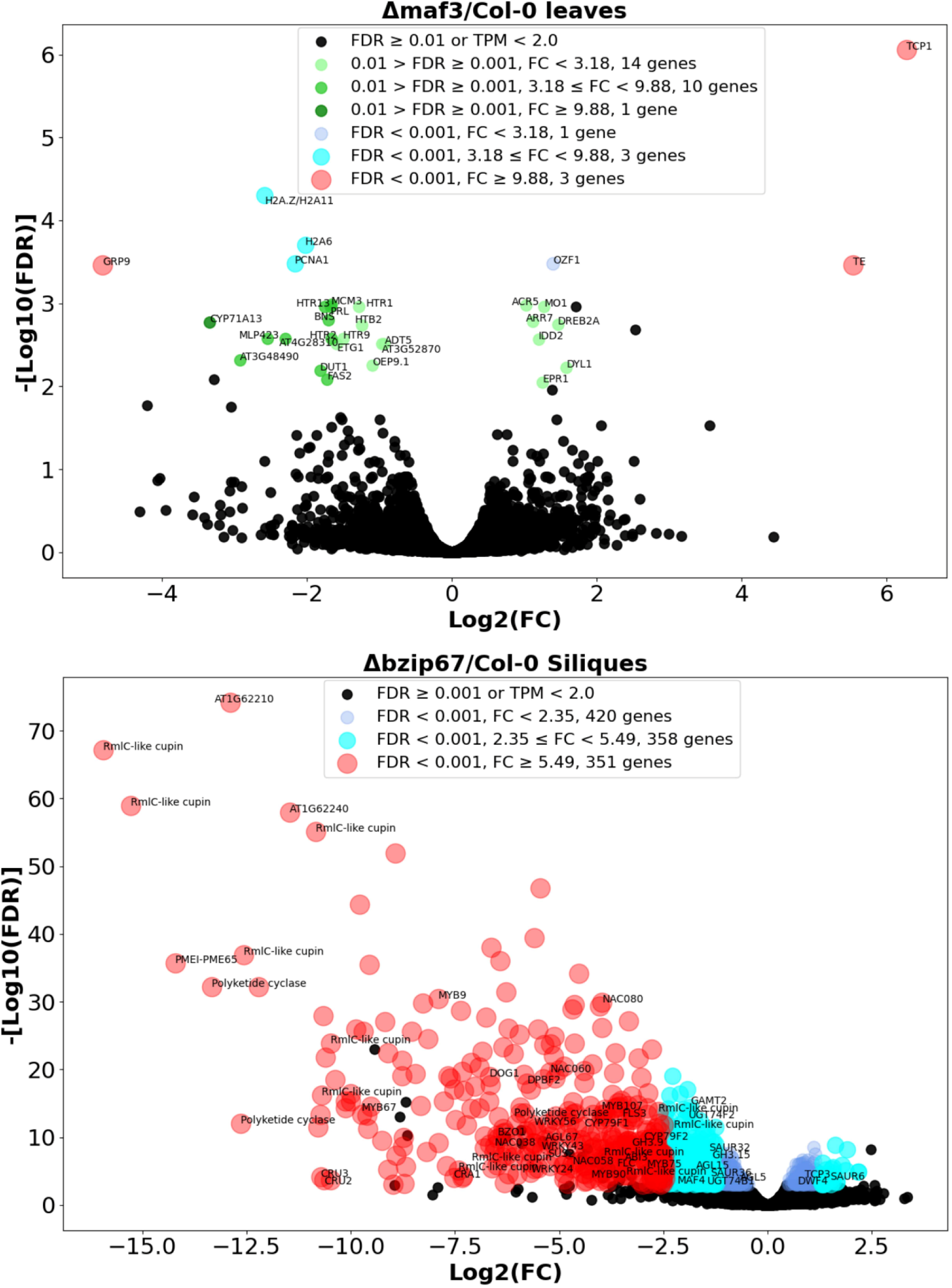
Differentially expressed *Arabidopsis* genes. Volcano plots of gene expression levels comparing wild-type and mutant lines of *Arabidopsis* at stage 4-6 as defined previously (Mizzotti *et al*., 2018) Fold-change (FC) thresholds were determined to have less than 5% (FC in MAF3 experiment samples ≥ 3.18, FC in BZIP67 experiment samples ≥ 2.35) or 1% (FC in MAF3 experiment samples ≥ 9.88, FC in BZIP67 experiment samples ≥ 5.49) likelihood to be surpassed by inter-replicate fold-changes, i.e., random fold-changes, as reported previously (Darbani & Stewart, 2014).

In contrast to *At*TCP18 (BRC1) and *At*TCP12 (BRC2), *At*TCP1 has possibly a role in induction of branching. TCP1 promotes the transcription of the brassinosteroid biosynthesis gene *Dwf*4 (Gao *et al*. 2015). Higher brassinosteroid levels can stabilize and accumulate the BRC1 suppressors BES1 and BZR1 in the nucleus (Yin *et al*. 2002). The *AtDwf*4 gene was up-regulated (1.7-fold) in *bzip*67 mutant plants (Fig. 5, Dataset S1). Similar to the phytoplasma triggered up-regulation of the *EpTcp*4 gene in poinsettia (Fig. 3a), *AtTcp*3 was upregulated (1.9-fold) in *Arabidopsis bzip*67 mutants when compared to the wild-type plants (Fig. 5, Dataset S1). Although TCP3 and TCP4 are not involved in axillary branch regulation, a role in shoot regeneration is however introduced (Yang *et al*. 2020). Another notable differential expression event was the 2.4-fold repression of *AtClamt* in the *bzip*67 mutant plants (Dataset S1). The gene *Clamt* is involved in the biosynthesis of methyl carlactonoate and when knocked-out, induces shoot branching through interference with strigolactone signaling in *Arabidopsis* (Mashiguchi *et al*. 2022).

We further investigated the *bzip*67 mutant mediated differentially regulated genes for DNA and protein level interactions. The *Arabidopsis* interaction viewer (Geisler-Lee *et al*. 2007; Dong *et al*. 2019 1) was used to extract experimentally confirmed interaction partners or targets for the differentially expressed genes (Fig. 6). Of most interest, we found *At*TCP3, 4, 8, 9, 10, 11, 14, 15, and 16 transcription factors and auxin transporters *At*PIN1, 2, 3, 4, 7, *At*DWARF14, *At*BES1, and many auxin and abscisic acid signaling proteins as protein-level interaction partners of the differentially expressed genes (Fig. 6a). In addition, the promoter region of more than half of the differentially expressed genes including *bZip*67 are found as potential targets for proteins coded by 28 of the differentially expressed genes (Fig. 6a, Fig. S7). There was no interacting partner for bZIP67 among the *bzip*67 mutant mediated differentially expressed genes. However, we found 18 other proteins interacting with bZIP67 (Fig. 6a). These interacting proteins can also interact with 13 proteins coded by the identified differentially expressed gene (Fig. S7). These indirect interactions were found to be enough to initiate all identified protein-DNA interactions at the next interaction cascade (Fig. S7). We applied the ForceAtlas2 graph algorithm for protein-protein network interaction analysis (Jacomy *et al*. 2014). By network analysis based on the interactions, we also noticed two distinct but still connected sub-clusters (Fig. 6b). One of the sub-clusters was found to be functionally involved in transmembrane transport and cell wall biogenesis and the other sub-cluster was found to be enriched in hormonal perception and signaling components (Fig. 6b).

**Fig. 6.**
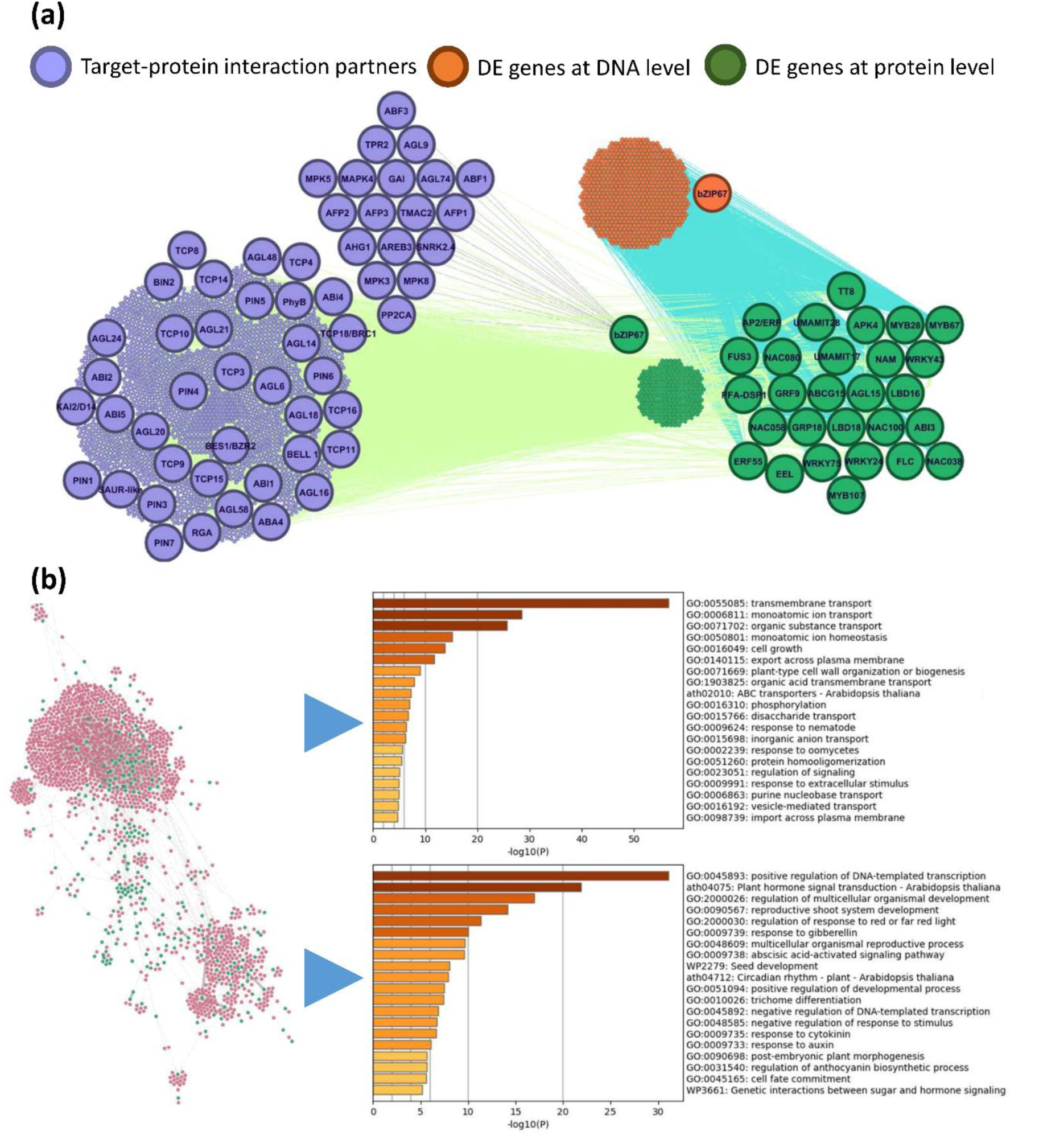
Protein-protein and protein-DNA interaction network for the differentially expressed genes and their interacting partners. (a) Protein-protein and protein-DNA interaction network. A selection of the interesting genes and proteins are enlarged and labeled. (b) Network analysis based on the interactions and pathway enrichment analysis for both of the identified interacting sub-clusters. Green nodes represent proteins coded by differentially expressed gene and pink nodes are interaction partners.

## Discussion

Shoot branching is an important commercial trait in poinsettia, which is usually obtained through phytoplasma infection, a tedious manual process that increases the risk of pathogen transmission and can introduce phenotypic variations (Lee *et al*. 2021). This study aimed at identifying candidate genes that could potentially be used in future breeding programs to produce stable branching lines of phytoplasma-free poinsettia through genetic manipulation (See Fig. 7). For this, we performed RNA sequencing and transcriptome analysis of phytoplasma-infected and -free poinsettia and identified several genes that were de-regulated after phytoplasma infection. Since the most dominant symptom of phytoplasma infection is free-branching, we hypothesized that these genes, particularly *Epmaf*3 and *EpbZip*67, were involved in branching as they were found to be highly repressed in the buds of infected poinsettia plants. When compared to wild-type plants, the *Arabidopsis maf*3 and *bzip*67 knock-out lines had more than two-fold higher number of primary branches, as was also the case in the *Arabidopsis Brc*1/*Tcp*18 knock-out mutant (Fig. 4).

**Fig. 7.**
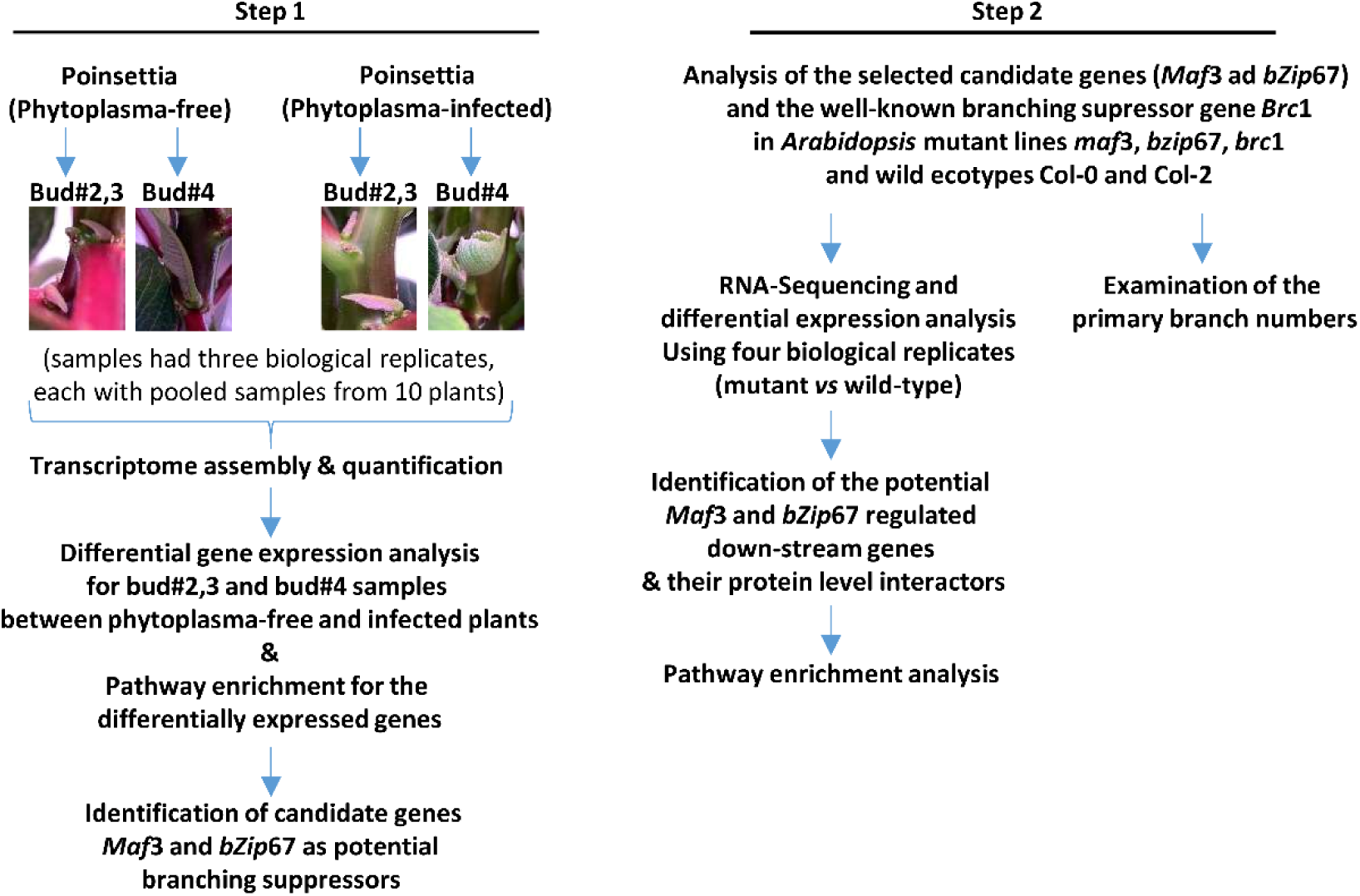
Experimental design of the study.

Transcriptome analyses in *Arabidopsis* were performed in leaves and siliques of *maf*3 and *bzip*67 mutant plants, respectively. This was a limitation and the reason to focus on the leaves and siliques and not on the buds was the impracticality of sampling sufficient synchronized tissue from extremely small *Arabidopsis* buds which could otherwise compromise the accuracy of transcriptome data. We found massive transcriptional repression in siliques of the *bzip*67 knock-out mutants, involving cellular transport and hormone signaling mechanisms. *AtClamt, AtGh*3.9, *AtGh*3.15*, AtSaur*32, *AtSaur*36, *AtAbi*3, and *AtGamt*2 were repressed while *AtTcp*3 and *AtDwf*4 were upregulated. The transcriptional attenuation bias of the genes which were differentially expressed in *bzip*67 mutant lines (Fig. 5, Dataset S1), is an indication of the fact that bZIP67 downstream signaling mainly involves transcriptional activations rather than suppression. MADS-box transcription factors such as MAF1, MAF5, AG, FLC, and SEP1 have been found as degradation targets of the phytoplasma effector protein SAP54 (MacLean *et al*. 2014; Kitazawa *et al*. 2022). Moreover, the *Maf*3 gene in *Euphorbia esula* (leafy spurge) has been found to induce crown bud endodormancy (Doğramaci *et al*. 2010; Doğramacı *et al*. 2011, 2013, 2014). In agreement with these reports, we here report the poinsettia *EpMaf*3 as the most highly repressed gene after phytoplasma infection in axillary buds. Indeed, T-DNA insertional disruption of this gene triggered shoot branching in *Arabidopsis*. We only found a handful of genes with transcriptional changes in leaves of the *maf*3 mutants, including *AtTcp*1 as the most up-regulated gene. Taken together, we introduce *EpMaf*3 and *EpbZip*67 as candidate genes regulating shoot branching and as promising targets for developing stable branching poinsettia lines by genetic engineering approaches. It is not clear whether *Maf*3 and *bZip*67 regulate branching upstream or downstream of *Brc*1/*Tcp*18 and *Brc*2/*Tcp*12 or functions independently from these TCPs. Nevertheless, the phytoplasma-mediated repression of *EpMaf*3 and *EpbZip*67 was accompanied by downregulation of *EpBrc*2/*Tcp*12 and upregulation of *EpTcp*4 in poinsettia. Furthermore, *AtTcp*1 was found to be upregulated in the leaves of the *Arabidopsis maf*3 knock-out line. While being a potential protein-level interaction partner for *At*TCP3, 4, 8, 9, 10, 11, 14, 15, 16, *At*PIN1, 2, 3, 4, 7, *At*DWARF14, and *At*BES1, *At*bZIP67 had also transcriptional regulatory effects on *AtDwf*4 and *AtTcp*3 genes in *Arabidopsis* siliques. It should be emphasized that findings from the *Arabidopsis* mutants alone have limited direct applicability to poinsettia. However, the revealed concordance between the results from the *Arabidopsis* experiments (i.e., the increased branching in the *Arabidopsis maf*3 and *bZip*67 knockout lines, along with the overlapping regulatory networks between well-known stem branching regulatory genes and the candidate genes *maf*3 and *bZip*67) and the poinsettia transcriptome analyses strengthens confidence in *maf*3 and *bZip*67 candidate genes for future genetic engineering efforts in poinsettia.

## Author Contributions

BD designed the experiments with help from others. BD performed the experiments and analyzed the data with help from CRI and IBH. MGM and JG were involved in growing and preparation of poinsettia plants and cuttings as well as project discussions. BD wrote the manuscript with help from coauthors. HBP and MN supervised the study and secured the funding.

## Data availability

RNA sequencing data is available under NCBI accession number PRJNA1171433.

## Supporting Information

Fig. S1 Quality check of total RNA samples before mRNA sequencing.

Fig. S2 Bud number 2, 3, and 4 from the top of the main shoot.

Fig. S3 Differentially expressed poinsettia genes with no homologues.

Fig. S4 T-DNA insertion sites and screening of the *Arabidopsis* mutant lines.

Fig. S5 Number of primary branches (PB) in the wild-type and mutant *Arabidopsis* thaliana lines.

Fig. S6 Data normalization for *Arabidopsis* transcriptome experiments.

Fig. S7 Protein-protein and protein-DNA interaction network for the differentially expressed genes and their interacting partners.

Fig. S8 Phylogenetic analysis for the *bZip*67 and *Maf*3 selected candidate genes.

Table S1 Sequencing statistics for poinsettia and *Arabidopsis* samples

Table S2 De novo assembly statistics of poinsettia transcriptome

## References

Aguilar-Martínez J.A., Poza-Carrión C., Cubas P. (2007) Arabidopsis BRANCHED1 acts as an integrator of branching signals within axillary buds. The Plant Cell 19:458–472.

Alonso J.M., Stepanova A.N., Leisse T.J., Kim C.J., Chen H., Shinn P., Stevenson D.K., Zimmerman J., Barajas P., Cheuk R., Gadrinab C., Heller C., Jeske A., Koesema E., Meyers C.C., Parker H., Prednis L., Ansari Y., Choy N., Deen H., Geralt M., Hazari N., Hom E., Karnes M., Mulholland C., Ndubaku R., Schmidt I., Guzman P., Aguilar-Henonin L., Schmid M., Weigel D., Carter D.E., Marchand T., Risseeuw E., Brogden D., Zeko A., Crosby W.L., Berry C.C., Ecker J.R. (2003) Genome-Wide Insertional Mutagenesis of Arabidopsis thaliana. Science 301:653–657.

Bai X., Correa V.R., Toruño T.Y., Ammar E.-D., Kamoun S., Hogenhout S.A. (2009) AY-WB phytoplasma secretes a protein that targets plant cell nuclei. Molecular plant-microbe interactions: MPMI 22:18–30.

Bai B., Zhang G., Pei B., Song Q., Hao X., Zhao L., Wu Y. (2023) The function of the phytoplasma effector SWP12 depends on the properties of two key amino acids. Journal of Biological Chemistry 299 [online] URL: https://www.jbc.org/article/S0021-9258(23)00184-9/abstract (accessed 26 September 2024).

Barbier F., Péron T., Lecerf M., Perez-Garcia M.-D., Barrière Q., Rolčík J., Boutet-Mercey S., Citerne S., Lemoine R., Porcheron B., Roman H., Leduc N., Le Gourrierec J., Bertheloot J., Sakr S. (2015) Sucrose is an early modulator of the key hormonal mechanisms controlling bud outgrowth in Rosa hybrida. Journal of Experimental Botany 66:2569–2582.

Beveridge C.A., Rameau C., Wijerathna-Yapa A. (2023) Lessons from a century of apical dominance research. Journal of Experimental Botany 74:3903–3922.

Bolger A.M., Lohse M., Usadel B. (2014) Trimmomatic: a flexible trimmer for Illumina sequence data. Bioinformatics 30:2114–2120.

Booker J., Auldridge M., Wills S., McCarty D., Klee H., Leyser O. (2004) MAX3/CCD7 is a carotenoid cleavage dioxygenase required for the synthesis of a novel plant signaling molecule. Current biology: CB 14:1232–1238.

Booker J., Sieberer T., Wright W., Williamson L., Willett B., Stirnberg P., Turnbull C., Srinivasan M., Goddard P., Leyser O. (2005) MAX1 Encodes a Cytochrome P450 Family Member that Acts Downstream of MAX3/4 to Produce a Carotenoid-Derived Branch-Inhibiting Hormone. Developmental Cell 8:443–449.

Boyes D.C., Zayed A.M., Ascenzi R., McCaskill A.J., Hoffman N.E., Davis K.R., Görlach J. (2001) Growth Stage –Based Phenotypic Analysis of Arabidopsis. The Plant Cell 13:1499–1510.

Bryant F.M., Hughes D., Hassani-Pak K., Eastmond P.J. (2019) Basic LEUCINE ZIPPER TRANSCRIPTION FACTOR67 Transactivates DELAY OF GERMINATION1 to Establish Primary Seed Dormancy in Arabidopsis. The Plant Cell 31:1276–1288.

Bryant D.M., Johnson K., DiTommaso T., Tickle T., Couger M.B., Payzin-Dogru D., Lee T.J., Leigh N.D., Kuo T.-H., Davis F.G., Bateman J., Bryant S., Guzikowski A.R., Tsai S.L., Coyne S., Ye W.W., Freeman R.M., Peshkin L., Tabin C.J., Regev A., Haas B.J., Whited J.L. (2017) A Tissue-Mapped Axolotl De Novo Transcriptome Enables Identification of Limb Regeneration Factors. Cell Reports 18:762–776.

Caicedo A.L., Richards C., Ehrenreich I.M., Purugganan M.D. (2009) Complex Rearrangements Lead to Novel Chimeric Gene Fusion Polymorphisms at the Arabidopsis thaliana MAF2-5 Flowering Time Gene Cluster. Molecular Biology and Evolution 26:699–711.

Chang S.H., Tan C.M., Wu C.-T., Lin T.-H., Jiang S.-Y., Liu R.-C., Tsai M.-C., Su L.-W., Yang J.-Y. (2018) Alterations of plant architecture and phase transition by the phytoplasma virulence factor SAP11. Journal of Experimental Botany 69:5389–5401.

Chen Y., Chen L., Lun A.T.L., Baldoni P.L., Smyth G.K. (2025) edgeR v4: powerful differential analysis of sequencing data with expanded functionality and improved support for small counts and larger datasets. Nucleic Acids Research 53:gkaf018.

Chen S., Song X., Zheng Q., Liu Y., Yu J., Zhou Y., Xia X. (2023) The transcription factor SPL13 mediates strigolactone suppression of shoot branching by inhibiting cytokinin synthesis in Solanum lycopersicum. Journal of Experimental Botany 74:5722–5735.

Christensen N.M., Nicolaisen M., Hansen M., Schulz A. (2004) Distribution of phytoplasmas in infected plants as revealed by real-time PCR and bioimaging. Molecular plant-microbe interactions: MPMI 17:1175–1184.

Czechowski T., Stitt M., Altmann T., Udvardi M.K., Scheible W.-R. (2005) Genome-Wide Identification and Testing of Superior Reference Genes for Transcript Normalization in Arabidopsis. Plant Physiology 139:5–17.

Danecek P., Bonfield J.K., Liddle J., Marshall J., Ohan V., Pollard M.O., Whitwham A., Keane T., McCarthy S.A., Davies R.M., Li H. (2021) Twelve years of SAMtools and BCFtools. GigaScience 10:giab008.

Darbani B., Noeparvar S., Borg S. (2015) Deciphering Mineral Homeostasis in Barley Seed Transfer Cells at Transcriptional Level. PloS One 10:e0141398.

Darbani B., Stewart C.N. (2014) Reproducibility and reliability assays of the gene expression-measurements. Journal of Biological Research-Thessaloniki 21:3.

Darbani B., Stewart C.N., Noeparvar S., Borg S. (2014) Correction of gene expression data: Performance-dependency on inter-replicate and inter-treatment biases. Journal of Biotechnology 188:100–109.

Dekkers B.J.W., Willems L., Bassel G.W., van Bolderen-Veldkamp R.P. (Marieke), Ligterink W., Hilhorst H.W.M., Bentsink L. (2012) Identification of Reference Genes for RT–qPCR Expression Analysis in Arabidopsis and Tomato Seeds. Plant and Cell Physiology 53:28–37.

Doebley J., Stec A., Gustus C. (1995) Teosinte Branched1 and the Origin of Maize: Evidence for Epistasis and the Evolution of Dominance. Genetics 141:333–346.

Doebley J., Stec A., Hubbard L. (1997) The evolution of apical dominance in maize. Nature 386:485–488.

Doğramaci M., Horvath D.P., Chao W.S., Foley M.E., Christoffers M.J., Anderson J.V. (2010) Low temperatures impact dormancy status, flowering competence, and transcript profiles in crown buds of leafy spurge. Plant Molecular Biology 73:207–226.

Doğramacı M., Foley M.E., Chao W.S., Christoffers M.J., Anderson J.V. (2013) Induction of endodormancy in crown buds of leafy spurge (Euphorbia esula L.) implicates a role for ethylene and cross-talk between photoperiod and temperature. Plant Molecular Biology 81:577–593.

Doğramacı M., Horvath D.P., Anderson J.V. (2014) Dehydration-induced endodormancy in crown buds of leafy spurge highlights involvement of MAF3- and RVE1-like homologs, and hormone signaling cross-talk. Plant Molecular Biology 86:409–424.

Doğramacı M., Horvath D.P., Christoffers M.J., Anderson J.V. (2011) Dehydration and vernalization treatments identify overlapping molecular networks impacting endodormancy maintenance in leafy spurge crown buds. Functional & Integrative Genomics 11:611–626.

Dong S., Lau V., Song R., Ierullo M., Esteban E., Wu Y., Sivieng T., Nahal H., Gaudinier A., Pasha A., Oughtred R., Dolinski K., Tyers M., Brady S.M., Grene R., Usadel B., Provart N.J. (2019) Proteome-wide, Structure-Based Prediction of Protein-Protein Interactions/New Molecular Interactions Viewer1[OPEN]. Plant Physiology 179:1893–1907.

Dong C., Zhang L., Zhang Q., Yang Y., Li D., Xie Z., Cui G., Chen Y., Wu L., Li Z., Liu G., Zhang X., Liu C., Chu J., Zhao G., Xia C., Jia J., Sun J., Kong X., Liu X. (2023) Tiller Number1 encodes an ankyrin repeat protein that controls tillering in bread wheat. Nature Communications 14:836.

Dröge-Laser W., Snoek B.L., Snel B., Weiste C. (2018) The Arabidopsis bZIP transcription factor family-an update. Current Opinion in Plant Biology 45:36–49.

Eddy S.R. (2011) Accelerated Profile HMM Searches. PLOS Computational Biology 7:e1002195.

Fang Z., Ji Y., Hu J., Guo R., Sun S., Wang X. (2020) Strigolactones and Brassinosteroids Antagonistically Regulate the Stability of the D53–OsBZR1 Complex to Determine *FC1* Expression in Rice Tillering. Molecular Plant 13:586–597.

Ferreira M.J., Silva J., Pinto S.C., Coimbra S. (2023) I Choose You: Selecting Accurate Reference Genes for qPCR Expression Analysis in Reproductive Tissues in Arabidopsis thaliana. Biomolecules 13:463.

Fichtner F., Barbier F.F., Annunziata M.G., Feil R., Olas J.J., Mueller-Roeber B., Stitt M., Beveridge C.A., Lunn J.E. (2021) Regulation of shoot branching in arabidopsis by trehalose 6-phosphate. New Phytologist 229:2135–2151.

Gao Y., Zhang D., Li J. (2015) TCP1 Modulates DWF4 Expression via Directly Interacting with the GGNCCC Motifs in the Promoter Region of DWF4 in Arabidopsis thaliana. Journal of Genetics and Genomics 42:383–392.

Geisler-Lee J., O’Toole N., Ammar R., Provart N.J., Millar A.H., Geisler M. (2007) A Predicted Interactome for Arabidopsis. Plant Physiology 145:317–329.

González-Grandío E., Pajoro A., Franco-Zorrilla J.M., Tarancón C., Immink R.G.H., Cubas P. (2017) Abscisic acid signaling is controlled by a BRANCHED1/HD-ZIP I cascade in Arabidopsis axillary buds. Proceedings of the National Academy of Sciences of the United States of America 114:E245–E254.

Grabherr M.G., Haas B.J., Yassour M., Levin J.Z., Thompson D.A., Amit I., Adiconis X., Fan L., Raychowdhury R., Zeng Q., Chen Z., Mauceli E., Hacohen N., Gnirke A., Rhind N., di Palma F., Birren B.W., Nusbaum C., Lindblad-Toh K., Friedman N., Regev A. (2011) Full-length transcriptome assembly from RNA-Seq data without a reference genome. Nature Biotechnology 29:644–652.

Gu X., Le C., Wang Y., Li Z., Jiang D., Wang Y., He Y. (2013) Arabidopsis FLC clade members form flowering-repressor complexes coordinating responses to endogenous and environmental cues. Nature Communications 4:1947.

Guindon S., Dufayard J.-F., Lefort V., Anisimova M., Hordijk W., Gascuel O. (2010) New Algorithms and Methods to Estimate Maximum-Likelihood Phylogenies: Assessing the Performance of PhyML 3.0. Systematic Biology 59:307–321.

Haas B.J., Papanicolaou A., Yassour M., Grabherr M., Blood P.D., Bowden J., Couger M.B., Eccles D., Li B., Lieber M., MacManes M.D., Ott M., Orvis J., Pochet N., Strozzi F., Weeks N., Westerman R., William T., Dewey C.N., Henschel R., LeDuc R.D., Friedman N., Regev A. (2013) De novo transcript sequence reconstruction from RNA-seq using the Trinity platform for reference generation and analysis. Nature Protocols 8:1494–1512.

Hamiaux C., Drummond R.S.M., Janssen B.J., Ledger S.E., Cooney J.M., Newcomb R.D., Snowden K.C. (2012) DAD2 is an α/β hydrolase likely to be involved in the perception of the plant branching hormone, strigolactone. Current biology: CB 22:2032–2036.

Hayward A., Stirnberg P., Beveridge C., Leyser O. (2009) Interactions between Auxin and Strigolactone in Shoot Branching Control. Plant Physiology 151:400–412.

Hoshi A., Oshima K., Kakizawa S., Ishii Y., Ozeki J., Hashimoto M., Komatsu K., Kagiwada S., Yamaji Y., Namba S. (2009) A unique virulence factor for proliferation and dwarfism in plants identified from a phytopathogenic bacterium. Proceedings of the National Academy of Sciences of the United States of America 106:6416–6421.

Hu J., Ji Y., Hu X., Sun S., Wang X. (2019) BES1 Functions as the Co-regulator of D53-like SMXLs to Inhibit BRC1 Expression in Strigolactone-Regulated Shoot Branching in Arabidopsis. Plant Communications 1:100014.

Jacomy M., Venturini T., Heymann S., Bastian M. (2014) ForceAtlas2, a Continuous Graph Layout Algorithm for Handy Network Visualization Designed for the Gephi Software. PLOS ONE 9:e98679.

Katyayini N.U., Rinne P.L.H., Tarkowská D., Strnad M., van der Schoot C. (2020) Dual Role of Gibberellin in Perennial Shoot Branching: Inhibition and Activation. Frontiers in Plant Science 11 [online] URL: https://www.frontiersin.org/journals/plant-science/articles/10.3389/fpls.2020.00736/full (accessed 23 July 2024).

Kitazawa Y., Iwabuchi N., Maejima K., Sasano M., Matsumoto O., Koinuma H., Tokuda R., Suzuki M., Oshima K., Namba S., Yamaji Y. (2022) A phytoplasma effector acts as a ubiquitin-like mediator between floral MADS-box proteins and proteasome shuttle proteins. The Plant Cell 34:1709–1723.

Kudo T., Sasaki Y., Terashima S., Matsuda-Imai N., Takano T., Saito M., Kanno M., Ozaki S., Suwabe K., Suzuki G., Watanabe M., Matsuoka M., Takayama S., Yano K. (2016) Identification of reference genes for quantitative expression analysis using large-scale RNA-seq data of *Arabidopsis thaliana* and model crop plants. Genes & Genetic Systems 91:111–125.

Langmead B., Salzberg S.L. (2012) Fast gapped-read alignment with Bowtie 2. Nature Methods 9:357–359.

Langmead B., Wilks C., Antonescu V., Charles R. (2019) Scaling read aligners to hundreds of threads on general-purpose processors. Bioinformatics 35:421–432.

Lee S., Chu C.-Y., Chu C.-C. (2021) Variability of Phytoplasma Infection Density in Poinsettia and Evaluation of its Association with the Level of Branching in Host Plants. Plant Disease 105:1539–1545.

Lee I.M., Klopmeyer M., Bartoszyk I.M., Gundersen-Rindal D.E., Chou T.S., Thomson K.L., Eisenreich R. (1997) Phytoplasma induced free-branching in commercial poinsettia cultivars. Nature Biotechnology 15:178–182.

Li B., Dewey C.N. (2011) RSEM: accurate transcript quantification from RNA-Seq data with or without a reference genome. BMC Bioinformatics 12:323.

Li H., Handsaker B., Wysoker A., Fennell T., Ruan J., Homer N., Marth G., Abecasis G., Durbin R., 1000 Genome Project Data Processing Subgroup (2009) The Sequence Alignment/Map format and SAMtools. Bioinformatics 25:2078–2079.

Lin H., Wang R., Qian Q., Yan M., Meng X., Fu Z., Yan C., Jiang B., Su Z., Li J., Wang Y. (2009) DWARF27, an Iron-Containing Protein Required for the Biosynthesis of Strigolactones, Regulates Rice Tiller Bud Outgrowth. The Plant Cell 21:1512–1525.

Linck H., Lankes C., Krüger E., Reineke A. (2019) Elimination of Phytoplasmas in Rubus Mother Plants by Tissue Culture Coupled with Heat Therapy. Plant Disease 103:1252–1255.

Lu Z., Yu H., Xiong G., Wang J., Jiao Y., Liu G., Jing Y., Meng X., Hu X., Qian Q., Fu X., Wang Y., Li J. (2013) Genome-Wide Binding Analysis of the Transcription Activator IDEAL PLANT ARCHITECTURE1 Reveals a Complex Network Regulating Rice Plant Architecture. The Plant Cell 25:3743–3759.

MacLean A.M., Orlovskis Z., Kowitwanich K., Zdziarska A.M., Angenent G.C., Immink R.G.H., Hogenhout S.A. (2014) Phytoplasma Effector SAP54 Hijacks Plant Reproduction by Degrading MADS-box Proteins and Promotes Insect Colonization in a RAD23-Dependent Manner. PLOS Biology 12:e1001835.

Manni M., Berkeley M.R., Seppey M., Zdobnov E.M. (2021) BUSCO: Assessing Genomic Data Quality and Beyond. Current Protocols 1:e323.

Martín-Trillo M., Cubas P. (2010) TCP genes: a family snapshot ten years later. Trends in Plant Science 15:31–39.

Mashiguchi K., Seto Y., Onozuka Y., Suzuki S., Takemoto K., Wang Y., Dong L., Asami K., Noda R., Kisugi T., Kitaoka N., Akiyama K., Bouwmeester H., Yamaguchi S. (2022) A carlactonoic acid methyltransferase that contributes to the inhibition of shoot branching in Arabidopsis. Proceedings of the National Academy of Sciences 119:e2111565119.

Mendes A., Kelly A.A., van Erp H., Shaw E., Powers S.J., Kurup S., Eastmond P.J. (2013) bZIP67 regulates the omega-3 fatty acid content of Arabidopsis seed oil by activating fatty acid desaturase3. The Plant Cell 25:3104–3116.

Minh B.Q., Nguyen M.A.T., von Haeseler A. (2013) Ultrafast Approximation for Phylogenetic Bootstrap. Molecular Biology and Evolution 30:1188–1195.

Mistry J., Chuguransky S., Williams L., Qureshi M., Salazar G.A., Sonnhammer E.L.L., Tosatto S.C.E., Paladin L., Raj S., Richardson L.J., Finn R.D., Bateman A. (2021) Pfam: The protein families database in 2021. Nucleic Acids Research 49:D412–D419.

Mizzotti C., Rotasperti L., Moretto M., Tadini L., Resentini F., Galliani B.M., Galbiati M., Engelen K., Pesaresi P., Masiero S. (2018) Time-Course Transcriptome Analysis of Arabidopsis Siliques Discloses Genes Essential for Fruit Development and Maturation. Plant Physiology 178:1249–1268.

Nakamura H., Xue Y.-L., Miyakawa T., Hou F., Qin H.-M., Fukui K., Shi X., Ito E., Ito S., Park S.-H., Miyauchi Y., Asano A., Totsuka N., Ueda T., Tanokura M., Asami T. (2013) Molecular mechanism of strigolactone perception by DWARF14. Nature Communications 4:2613.

Nguyen L.-T., Schmidt H.A., von Haeseler A., Minh B.Q. (2015) IQ-TREE: A Fast and Effective Stochastic Algorithm for Estimating Maximum-Likelihood Phylogenies. Molecular Biology and Evolution 32:268–274.

Ni J., Gao C., Chen M.-S., Pan B.-Z., Ye K., Xu Z.-F. (2015) Gibberellin Promotes Shoot Branching in the Perennial Woody Plant Jatropha curcas. Plant and Cell Physiology 56:1655–1666.

Nicolas M., Cubas P. (2016) TCP factors: new kids on the signaling block. Current Opinion in Plant Biology 33:33–41.

Nitarska D., Boehm R., Debener T., Lucaciu R.C., Halbwirth H. (2021) First genome edited poinsettias: targeted mutagenesis of flavonoid 3ʹ-hydroxylase using CRISPR/Cas9 results in a colour shift. Plant Cell, Tissue and Organ Culture 147:49–60.

Oshima K., Maejima K., Isobe Y., Endo A., Namba S., Yamaji Y. (2023) Molecular mechanisms of plant manipulation by secreting effectors of phytoplasmas. Physiological and Molecular Plant Pathology 125:102009.

Patil S.B., Barbier F.F., Zhao J., Zafar S.A., Uzair M., Sun Y., Fang J., Perez-Garcia M.-D., Bertheloot J., Sakr S., Fichtner F., Chabikwa T.G., Yuan S., Beveridge C.A., Li X. (2022) Sucrose promotes D53 accumulation and tillering in rice. New Phytologist 234:122–136.

Pecher P., Moro G., Canale M.C., Capdevielle S., Singh A., MacLean A., Sugio A., Kuo C.-H., Lopes J.R.S., Hogenhout S.A. (2019) Phytoplasma SAP11 effector destabilization of TCP transcription factors differentially impact development and defence of Arabidopsis versus maize. PLoS Pathogens 15:e1008035.

Rieu I., Eriksson S., Powers S.J., Gong F., Griffiths J., Woolley L., Benlloch R., Nilsson O., Thomas S.G., Hedden P., Phillips A.L. (2008) Genetic Analysis Reveals That C19-GA 2-Oxidation Is a Major Gibberellin Inactivation Pathway in Arabidopsis. The Plant Cell 20:2420–2436.

Robinson M.D., McCarthy D.J., Smyth G.K. (2010) edgeR: a Bioconductor package for differential expression analysis of digital gene expression data. Bioinformatics 26:139–140.

Roesler K., Lu C., Thomas J., Xu Q., Vance P., Hou Z., Williams R.W., Liu L., Owens M.A., Habben J.E. (2021) Arabidopsis Carboxylesterase 20 Binds Strigolactone and Increases Branches and Tillers When Ectopically Expressed in Arabidopsis and Maize. Frontiers in Plant Science 12:639401.

Sakamoto T., Ortega J.M. (2021) Taxallnomy: an extension of NCBI Taxonomy that produces a hierarchically complete taxonomic tree. BMC Bioinformatics 22:388.

Salam B.B., Barbier F., Danieli R., Teper-Bamnolker P., Ziv C., Spíchal L., Aruchamy K., Shnaider Y., Leibman D., Shaya F., Carmeli-Weissberg M., Gal-On A., Jiang J., Ori N., Beveridge C., Eshel D. (2021) Sucrose promotes stem branching through cytokinin. Plant Physiology 185:1708–1721.

Schwartz S.H., Qin X., Loewen M.C. (2004) The biochemical characterization of two carotenoid cleavage enzymes from Arabidopsis indicates that a carotenoid-derived compound inhibits lateral branching. The Journal of Biological Chemistry 279:46940–46945.

Shen J., Zhang Y., Ge D., Wang Z., Song W., Gu R., Che G., Cheng Z., Liu R., Zhang X. (2019) CsBRC1 inhibits axillary bud outgrowth by directly repressing the auxin efflux carrier CsPIN3 in cucumber. Proceedings of the National Academy of Sciences of the United States of America 116:17105–17114.

Sorefan K., Booker J., Haurogné K., Goussot M., Bainbridge K., Foo E., Chatfield S., Ward S., Beveridge C., Rameau C., Leyser O. (2003) MAX4 and RMS1 are orthologous dioxygenase-like genes that regulate shoot branching in Arabidopsis and pea. Genes & Development 17:1469–1474.

Stirnberg P., Furner I.J., Ottoline Leyser H.M. (2007) MAX2 participates in an SCF complex which acts locally at the node to suppress shoot branching. The Plant Journal: For Cell and Molecular Biology 50:80–94.

Stirnberg P., van de Sande K., Leyser H.M.O. (2002) MAX1 and MAX2 control shoot lateral branching in Arabidopsis. Development 129:1131–1141.

Sugio A., Kingdom H.N., MacLean A.M., Grieve V.M., Hogenhout S.A. (2011) Phytoplasma protein effector SAP11 enhances insect vector reproduction by manipulating plant development and defense hormone biosynthesis. Proceedings of the National Academy of Sciences of the United States of America 108:E1254–1263.

Tamura K., Stecher G., Kumar S. (2021) MEGA11: Molecular Evolutionary Genetics Analysis Version 11. Molecular Biology and Evolution 38:3022–3027.

Tatematsu K., Ward S., Leyser O., Kamiya Y., Nambara E. (2005) Identification of cis-Elements That Regulate Gene Expression during Initiation of Axillary Bud Outgrowth in Arabidopsis. Plant Physiology 138:757–766.

Tegenfeldt F., Kuznetsov D., Manni M., Berkeley M., Zdobnov E.M., Kriventseva E.V. (2025) OrthoDB and BUSCO update: annotation of orthologs with wider sampling of genomes. Nucleic Acids Research 53:D516–D522.

The UniProt Consortium (2025) UniProt: the Universal Protein Knowledgebase in 2025. Nucleic Acids Research 53:D609–D617.

Vilperte V., Lucaciu C.R., Halbwirth H., Boehm R., Rattei T., Debener T. (2019) Hybrid de novo transcriptome assembly of poinsettia (Euphorbia pulcherrima Willd. Ex Klotsch) bracts. BMC Genomics 20:900.

Waese J., Fan J., Pasha A., Yu H., Fucile G., Shi R., Cumming M., Kelley L.A., Sternberg M.J., Krishnakumar V., Ferlanti E., Miller J., Town C., Stuerzlinger W., Provart N.J. (2017) ePlant: Visualizing and Exploring Multiple Levels of Data for Hypothesis Generation in Plant Biology. The Plant Cell 29:1806–1821.

Wang S., Shi Y., Zhou Y., Hu W., Liu F. (2024) Full-length transcriptome sequencing of Arabidopsis plants provided new insights into the autophagic regulation of photosynthesis. Scientific Reports 14:14588.

Wang B., Smith S.M., Li J. (2018) Genetic Regulation of Shoot Architecture. Annual Review of Plant Biology 69:437–468.

Wang Y., Sun S., Zhu W., Jia K., Yang H., Wang X. (2013) Strigolactone/MAX2-induced degradation of brassinosteroid transcriptional effector BES1 regulates shoot branching. Developmental Cell 27:681–688.

Woody S.T., Austin-Phillips S., Amasino R.M., Krysan P.J. (2007) The WiscDsLox T-DNA collection: an arabidopsis community resource generated by using an improved high-throughput T-DNA sequencing pipeline. Journal of Plant Research 120:157–165.

Xu E., Chai L., Zhang S., Yu R., Zhang X., Xu C., Hu Y. (2021) Catabolism of strigolactones by a carboxylesterase. Nature Plants 7:1495–1504.

Yang W., Choi M.-H., Noh B., Noh Y.-S. (2020) De Novo Shoot Regeneration Controlled by HEN1 and TCP3/4 in Arabidopsis. Plant & Cell Physiology 61:1600–1613.

Yin Y., Wang Z.Y., Mora-Garcia S., Li J., Yoshida S., Asami T., Chory J. (2002) BES1 accumulates in the nucleus in response to brassinosteroids to regulate gene expression and promote stem elongation. Cell 109:181–191.

Yu Z., Zhang F., Friml J., Ding Z. (2022) Auxin signaling: Research advances over the past 30 years. Journal of Integrative Plant Biology 64:371–392.

Zhang S., Li R., Zhang L., Chen S., Xie M., Yang L., Xia Y., Foyer C.H., Zhao Z., Lam H.-M. (2020) New insights into Arabidopsis transcriptome complexity revealed by direct sequencing of native RNAs. Nucleic Acids Research 48:7700–7711.

Zheng Z., Guo Y., Novák O., Chen W., Ljung K., Noel J.P., Chory J. (2016) Local auxin metabolism regulates environment-induced hypocotyl elongation. Nature Plants 2:1–9.

Zhou Y., Zhou B., Pache L., Chang M., Khodabakhshi A.H., Tanaseichuk O., Benner C., Chanda S.K. (2019) Metascape provides a biologist-oriented resource for the analysis of systems-level datasets. Nature Communications 10:1–10.

