## Supplementary information for "Phytoplasma mediated transcriptional changes in poinsettia buds suggest MAF3 and bZIP67 transcription factors as potential suppressors of shoot branching"

Authors: Behrooz Darbani[1#*], Christina R. Ingvardsen[1], Inger B. Holme[1], Marianne Gellert Møller[2], Jacob Graff[2], Henrik Brinch-Pedersen[1], Mogens Nicolaisen[1*]
[1]Department of Agroecology, Faculty of Technical Sciences, Aarhus University, DK-4200 Slagelse, Denmark

[1]Department of Agroecology, Faculty of Technical Sciences, Aarhus University, DK-4200 Slagelse, Denmark
[2]Graff Breeding A/S, Viborgvej 717, 8471 Sabro, Denmark
[#]Present address: Department of Clinical Research, Hospital Sønderjylland, Aabenraa, Denmark, Department of Regional Health Research, University of Southern Denmark, Aabenraa, Denmark

The following Supporting Information is available for this article:

**Fig. S1 Quality check of total RNA samples before mRNA sequencing.**

**Fig. S2 Bud number 2, 3, and 4 from the top of the main shoot.**

**Fig. S3 Differentially expressed poinsettia genes with no homologues.**

**Fig. S4 T-DNA insertion sites and screening of the Arabidopsis mutant lines.**

**Fig. S5 Number of primary branches (PB) in the wild-type and mutant *Arabidopsis thaliana* lines.**

**Fig. S6 Data normalization for Arabidopsis transcriptome experiments.**

**Fig. S7 Protein-protein and protein-DNA interaction network for the differentially expressed genes and their interacting partners.**

**Fig. S8 Phylogenetic analysis for the bZip67 and Maf3 selected candidate genes.**

**Table S1 Sequencing statistics for poinsettia and Arabidopsis samples**

**Table S2 De novo assembly statistics of poinsettia transcriptome**

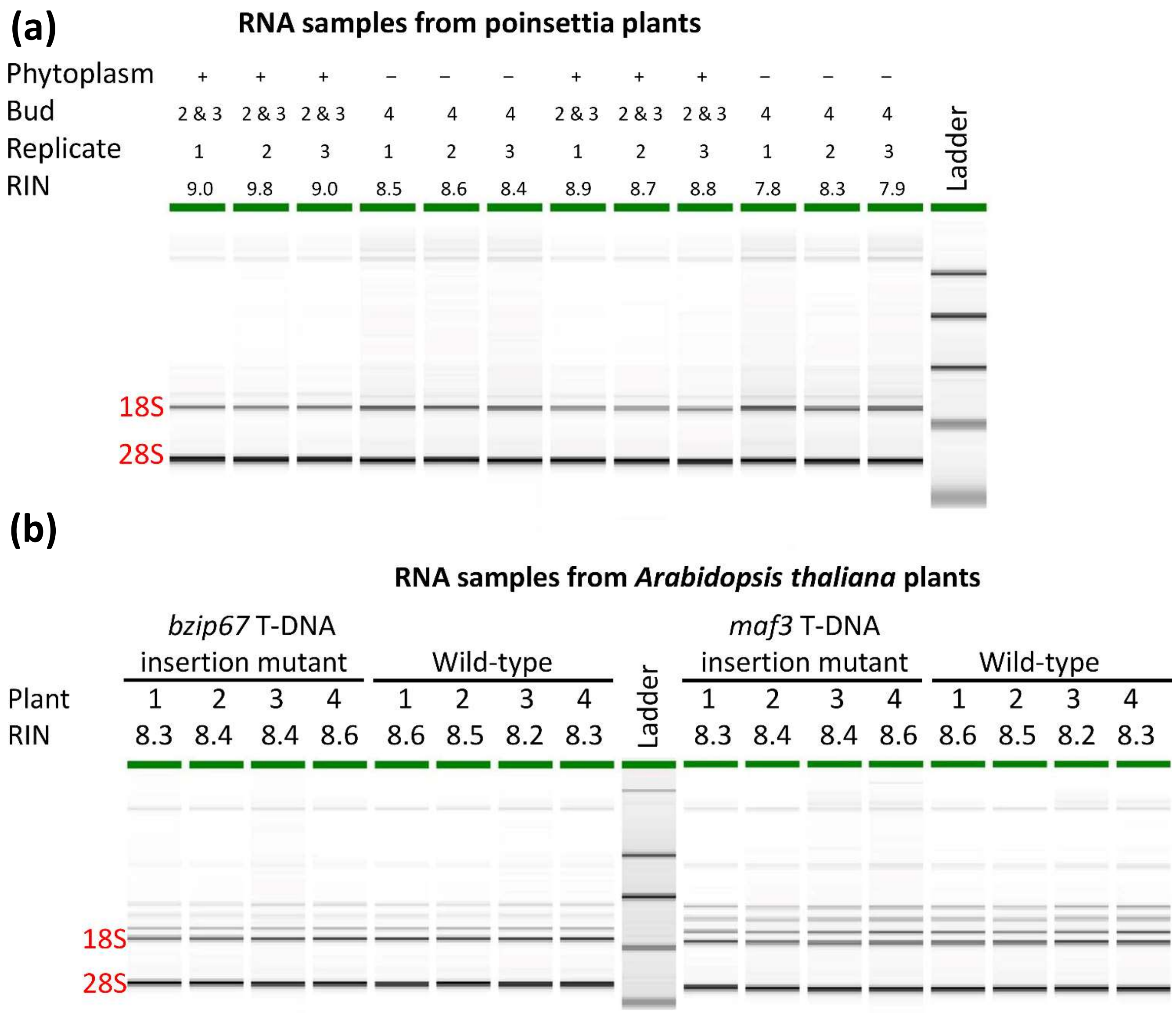


**Fig. S1 Quality check of total RNA samples before mRNA sequencing.** The RNA integrity number (RIN) varies between 1 (very degraded) and 10 (minimum degradation) estimated by Agilent 2100 Bioanalyzer System and its associated RNA 6000 Nano assay kit. Each replicate was a pool of 10 plants.

Phytoplasma-free
Bud no. 2 & 3

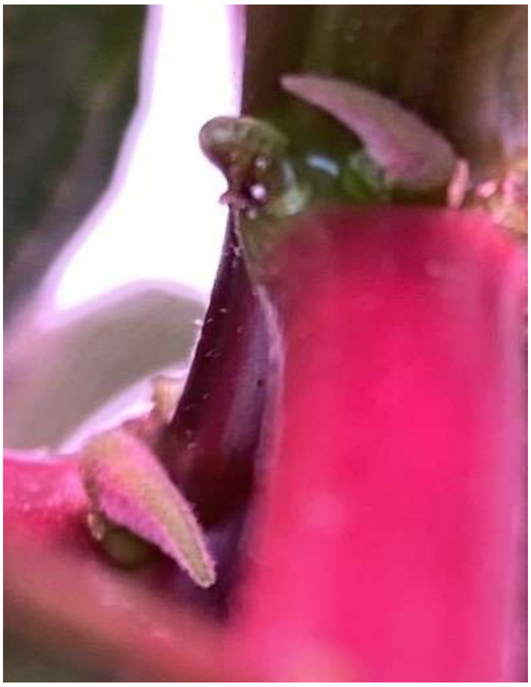

Phytoplasma-infected
Bud no. 2 & 3

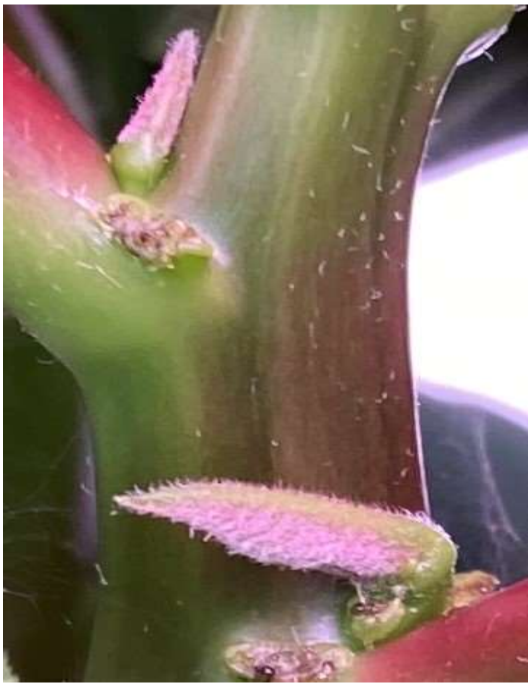

Phytoplasma-free
Bud no. 4

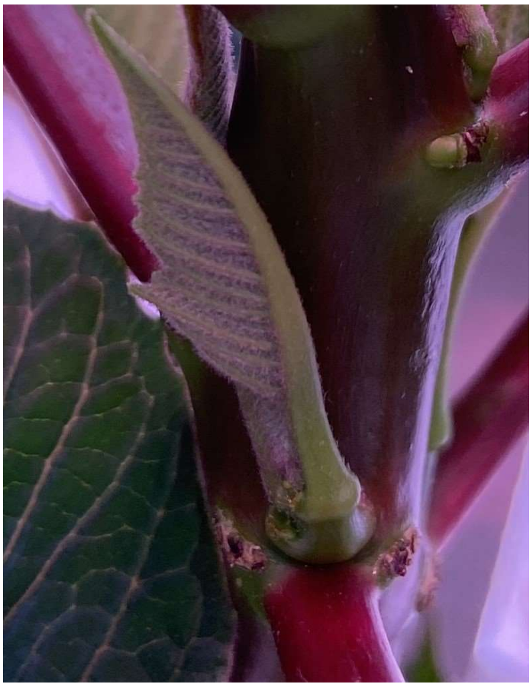

Phytoplasma-infected
Bud no. 4

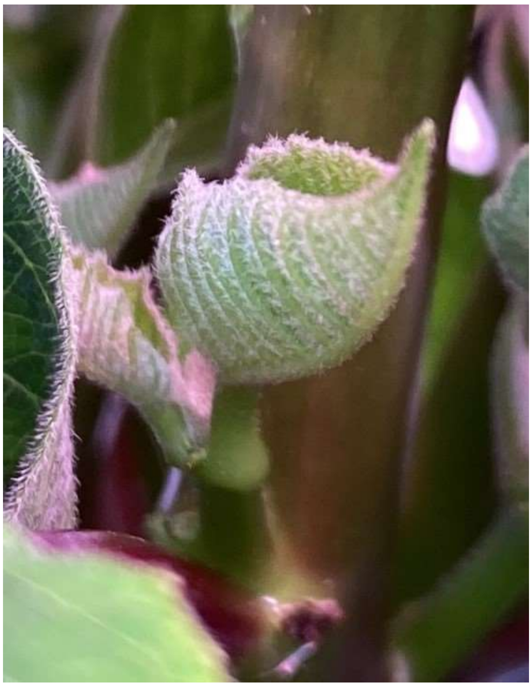

**Fig. S2 Bud number 2, 3, and 4 from the top of the main shoot.**

(a)

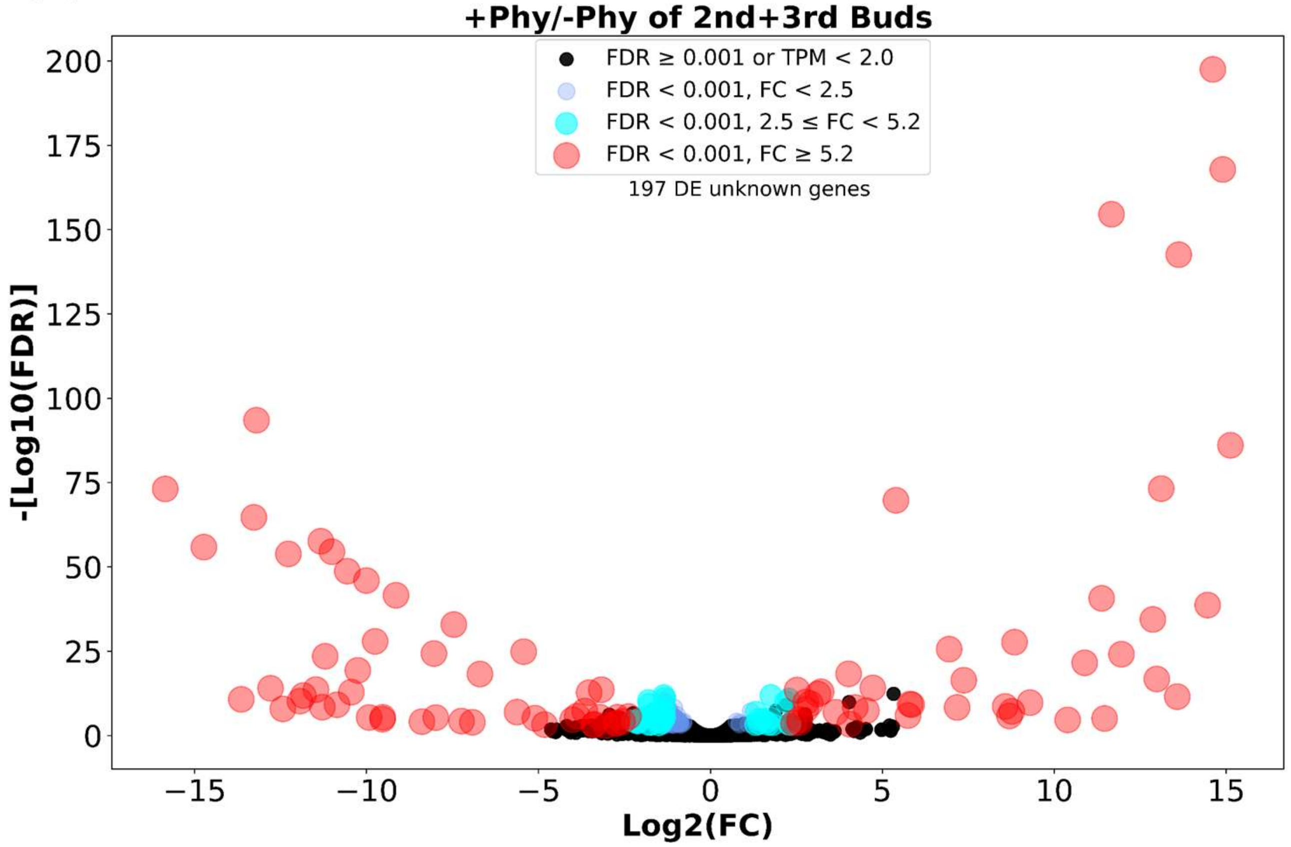


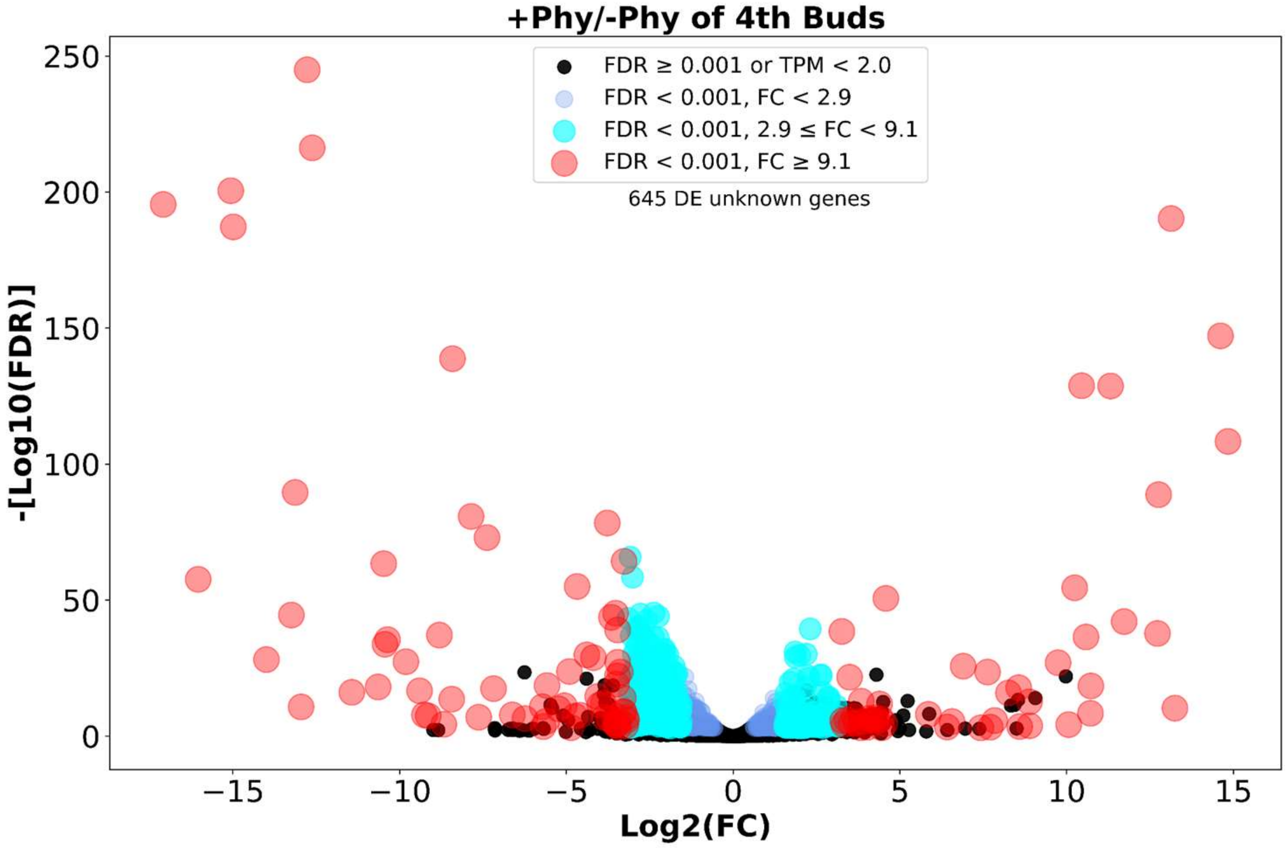


**Fig. S3 continued.**

(b)

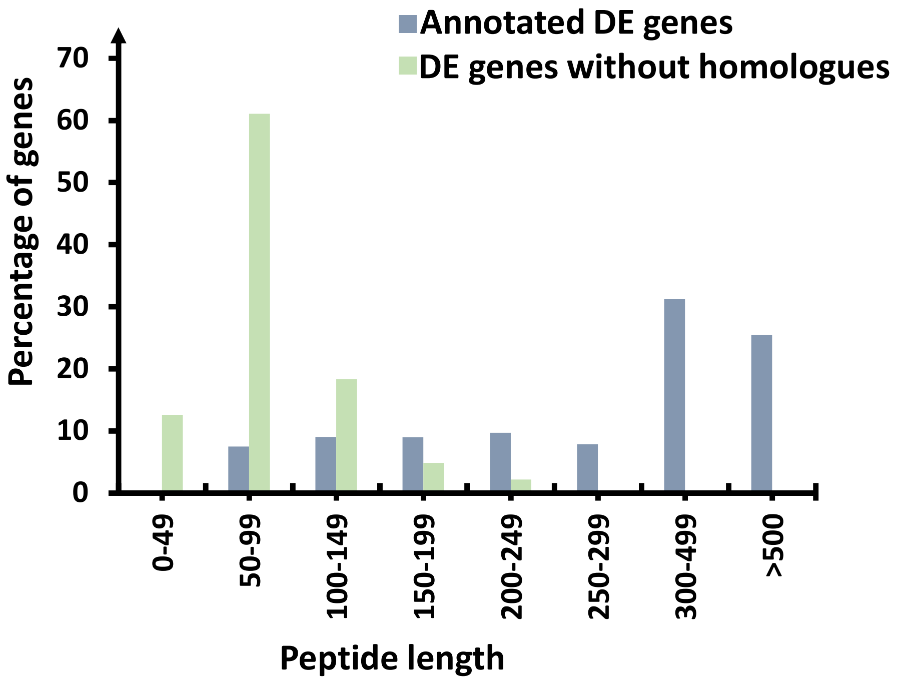


**Fig. S3 Differentially expressed poinsettia genes with no homologues.** (a) Volcano plots of gene expression levels when comparing phytoplasma-infected and phytoplasma-free poinsettia plants. Fold-change (FC) thresholds were determined to have less than 5% (FC for 2$^{nd}$+3$^{rd}$ bud samples ≥ 2.5, FC for 4$^{th}$ bud samples ≥ 2.9) or 1% (FC for 2$^{nd}$+3$^{rd}$ bud samples ≥ 5.2, FC for 4$^{th}$ bud samples ≥ 9.1) likelihood to be surpassed by inter-replicate fold-changes, i.e., random fold-changes, as explained in Material and Methods. (b) Length distribution of the proteins coded by the differentially expressed (DE) genes.

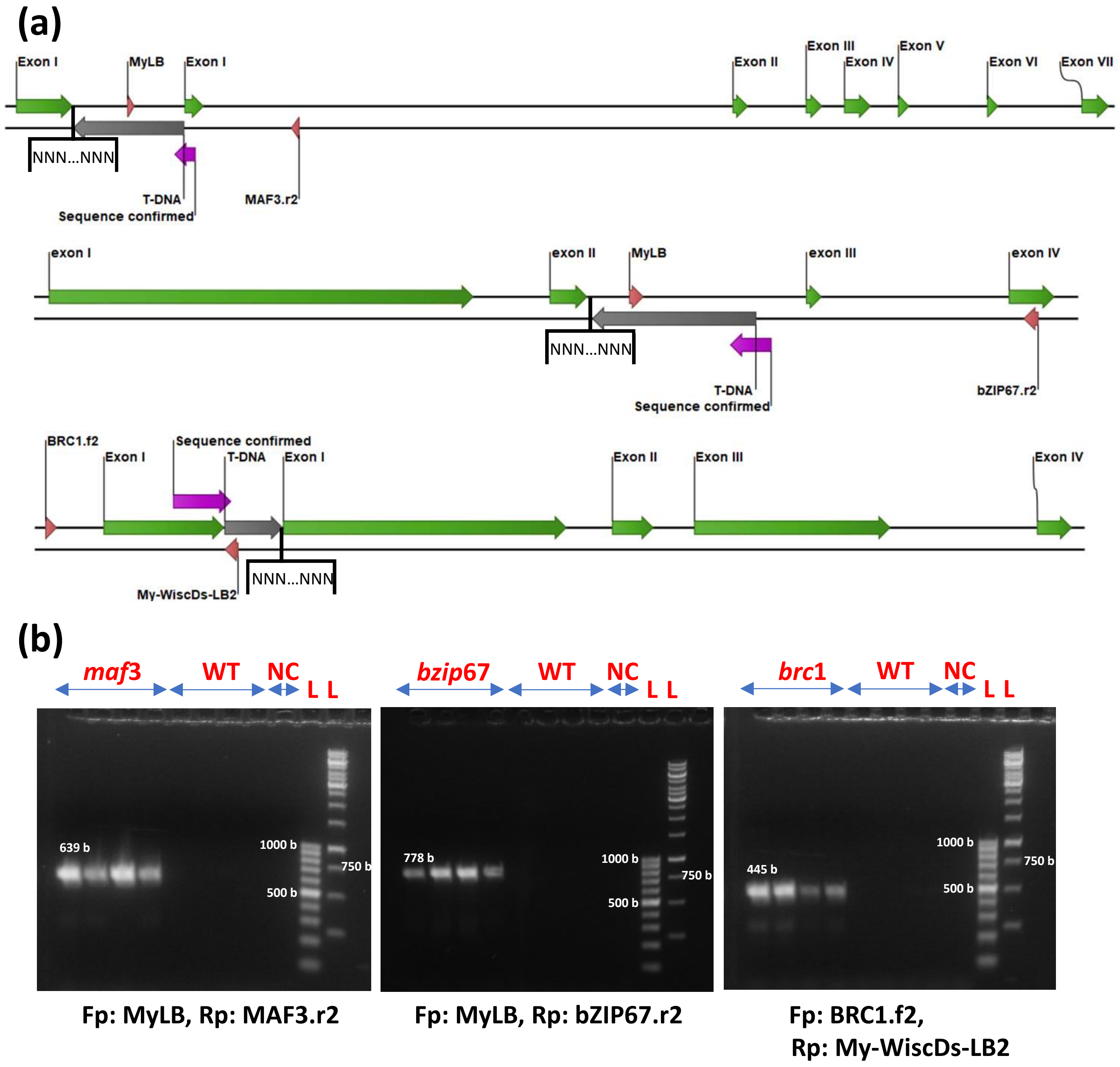


**Fig. S4 T-DNA insertion sites and screening of the *Arabidopsis* mutant lines.** (a) Gene structure illustration together with T-DNA insertion sites and screening primers. MyLB, MAF3.r2, bZIP67.r2, BRC1.f2, and My-WiscDs-LB2 are screening primers. Upper panel: *Maf*3, Middle panel: *bZip*67, bottom panel: *Brc*1. (b) PCR screening of the *Arabidopsis* wildtype lines (Col-0 [NASC ID: N70000] or Col-2 [NASC ID: N907]) and three T-DNA insertion mutant lines *maf*3 [NASC ID: N674795], *bzip*67 [NASC ID: N666597], and *brc*1[NASC ID: N857231]. $F_P$ and $R_P$ are forward and reverse primers. NC: Negative control, L: ladder.

(a)

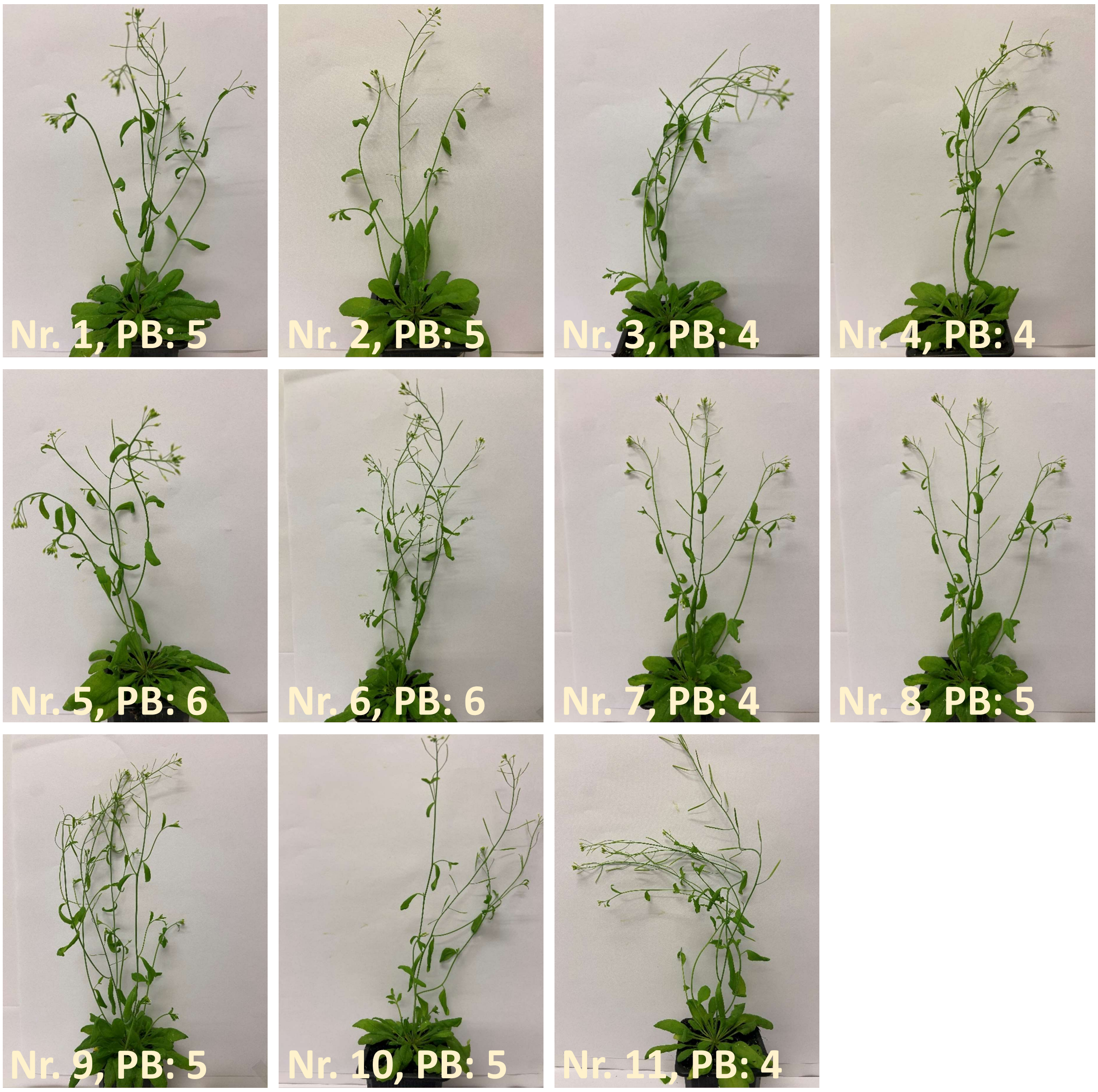


(b)

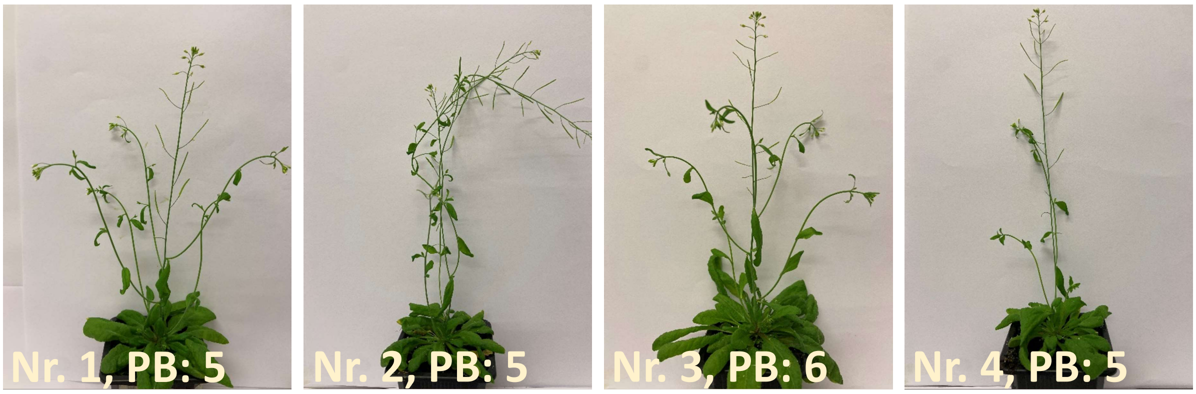


**Fig. S5 continued.**

**(b)**

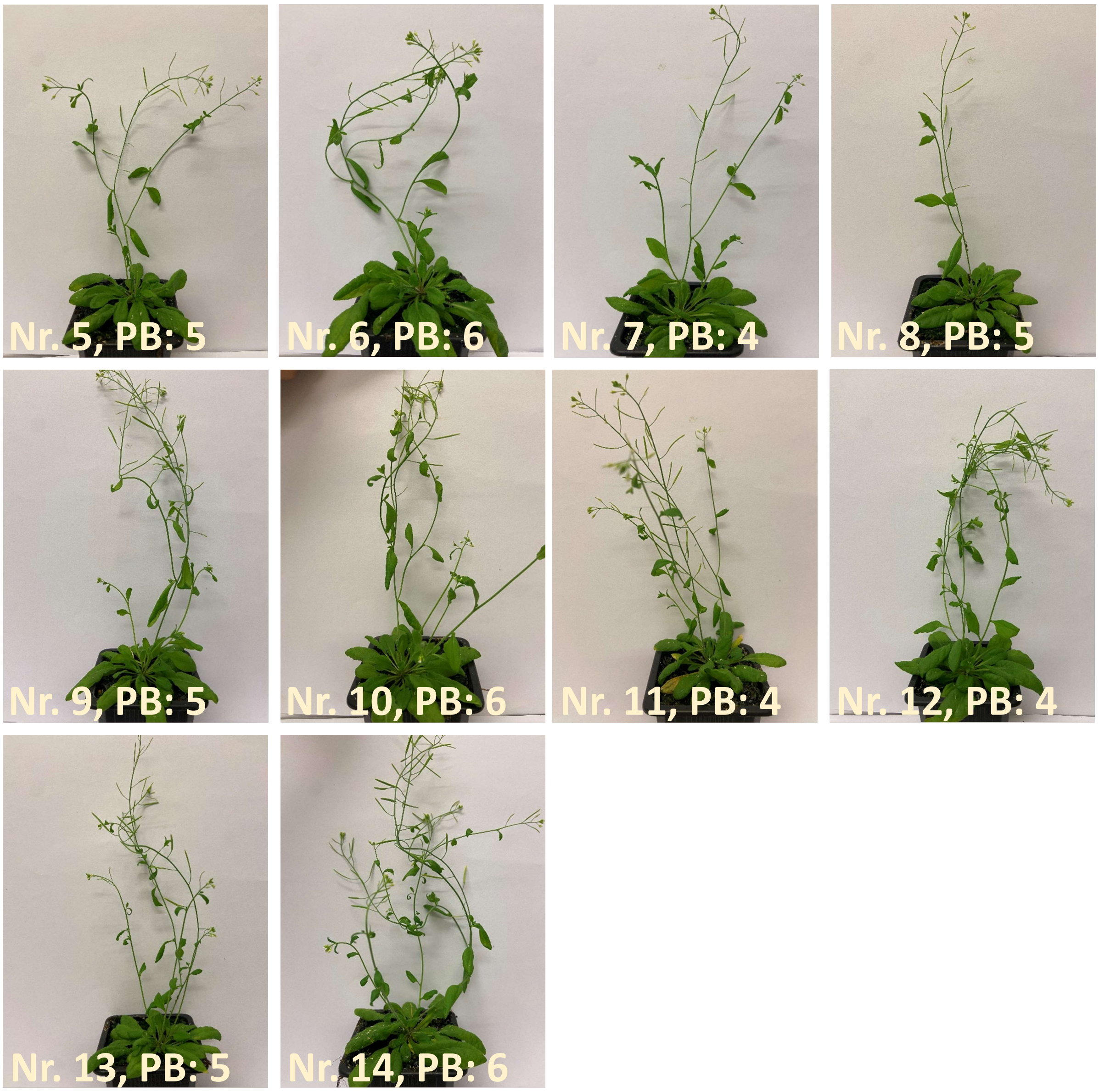


**(c)**

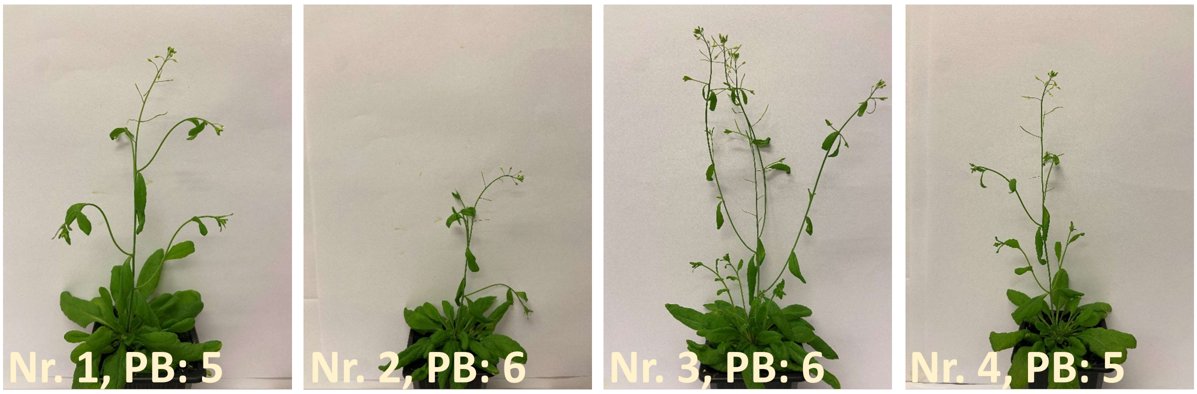


**Fig. S5 continued.**

(c)

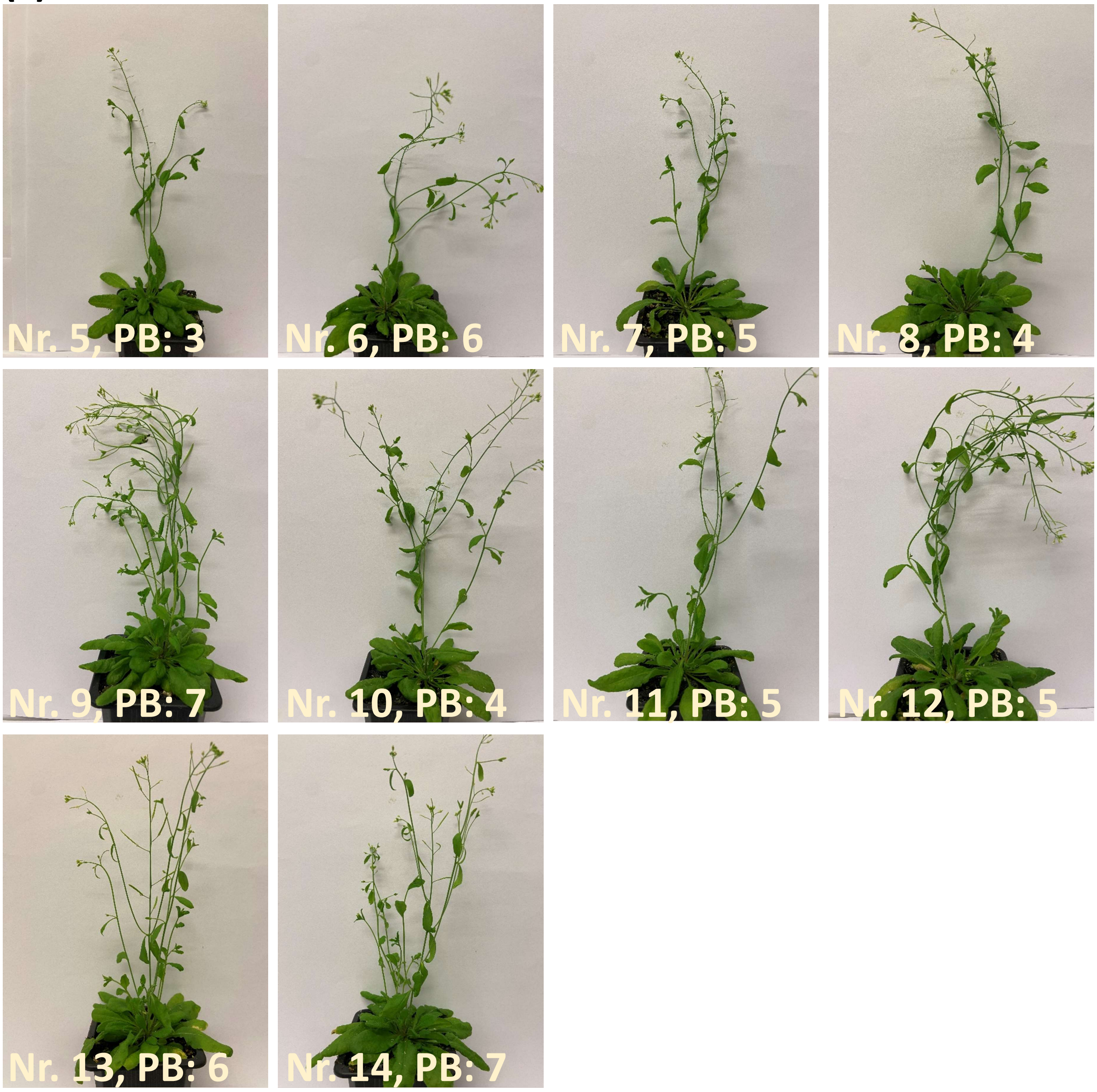


(d)

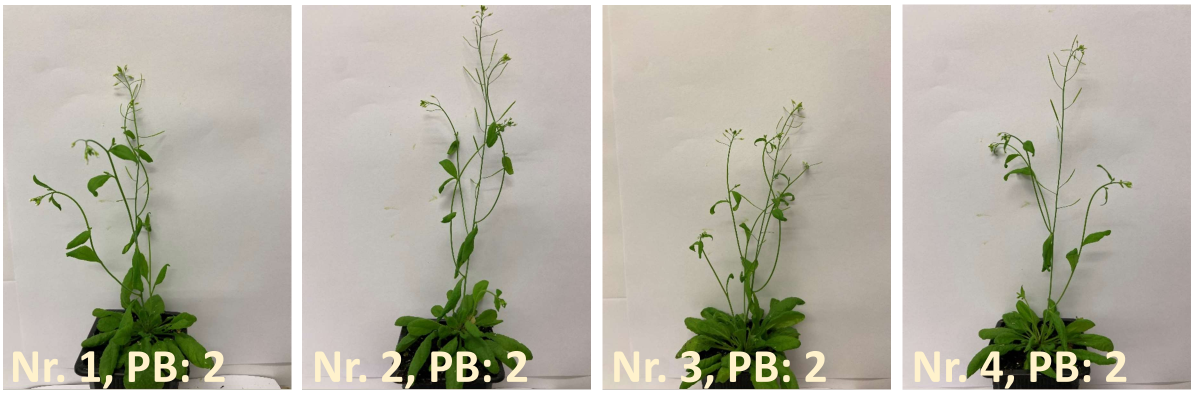


**Fig. S5 continued.**

(d)

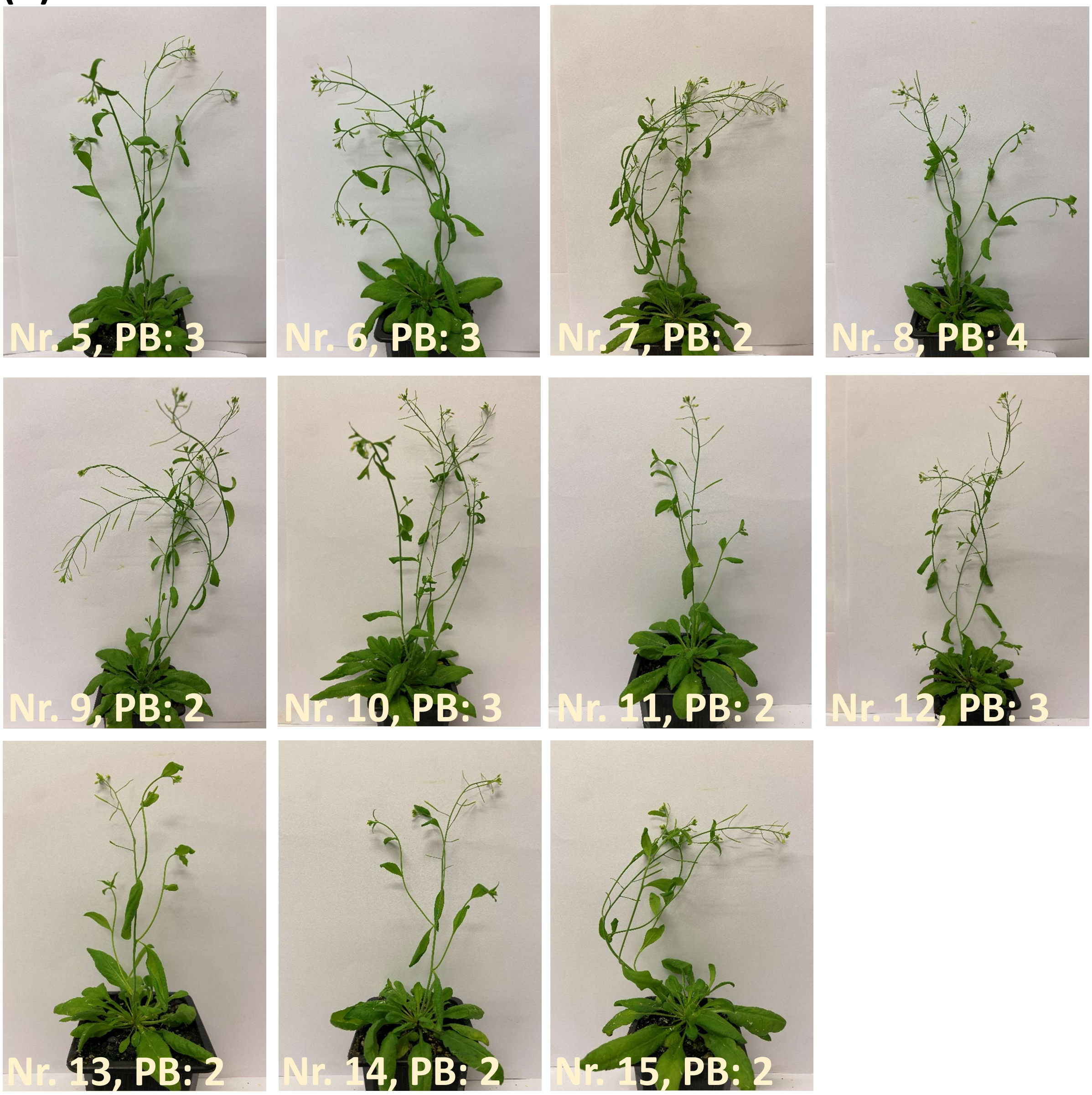


(e)

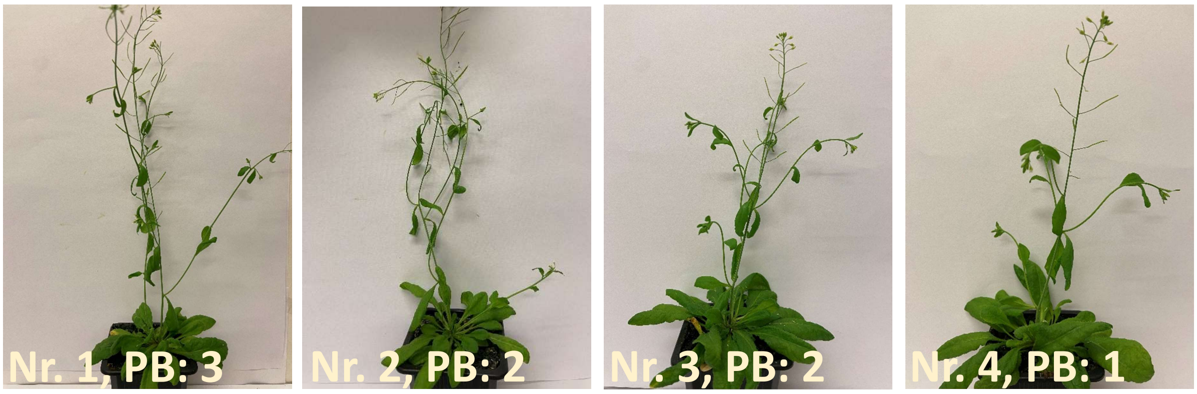


**Fig. S5 continued.**

**(e)**

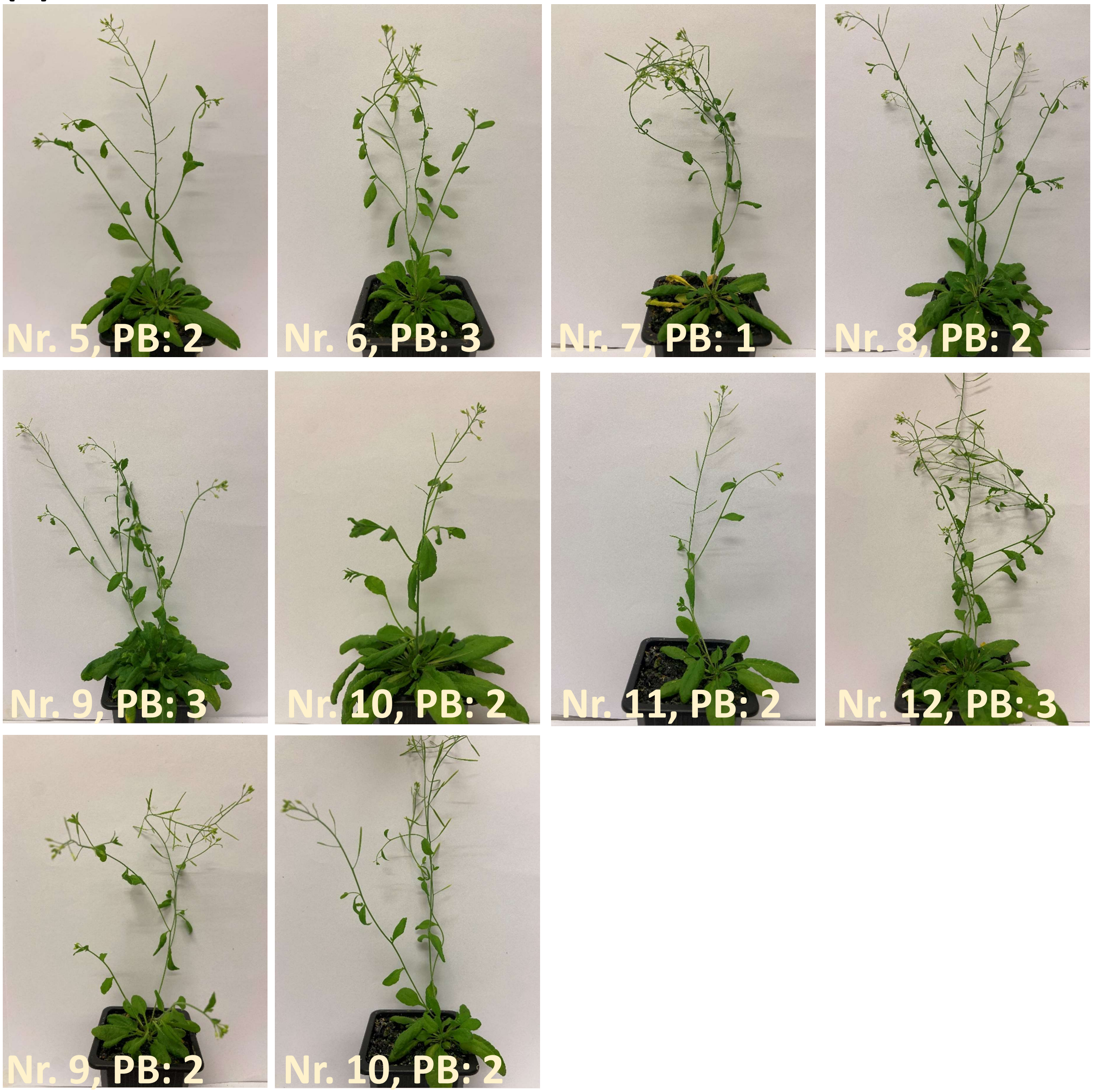


**Fig. S5 Number of primary branches (PB) in the wild-type and mutant *Arabidopsis thaliana* lines.** The number of primary branches (stems) for three T-DNA insertion mutant line (a) *maf*3 [NASC ID: N674795], (b) *bzip*67 [NASC ID: N666597], (c) *brc*1 [NASC ID: N857231]), (d) wild-type Col-0 [NASC ID: N70000] and (e) wild-type Col-2 [NASC ID: N907]). The number of primary branches (stems) were examined two weeks after inflorescence emergence.

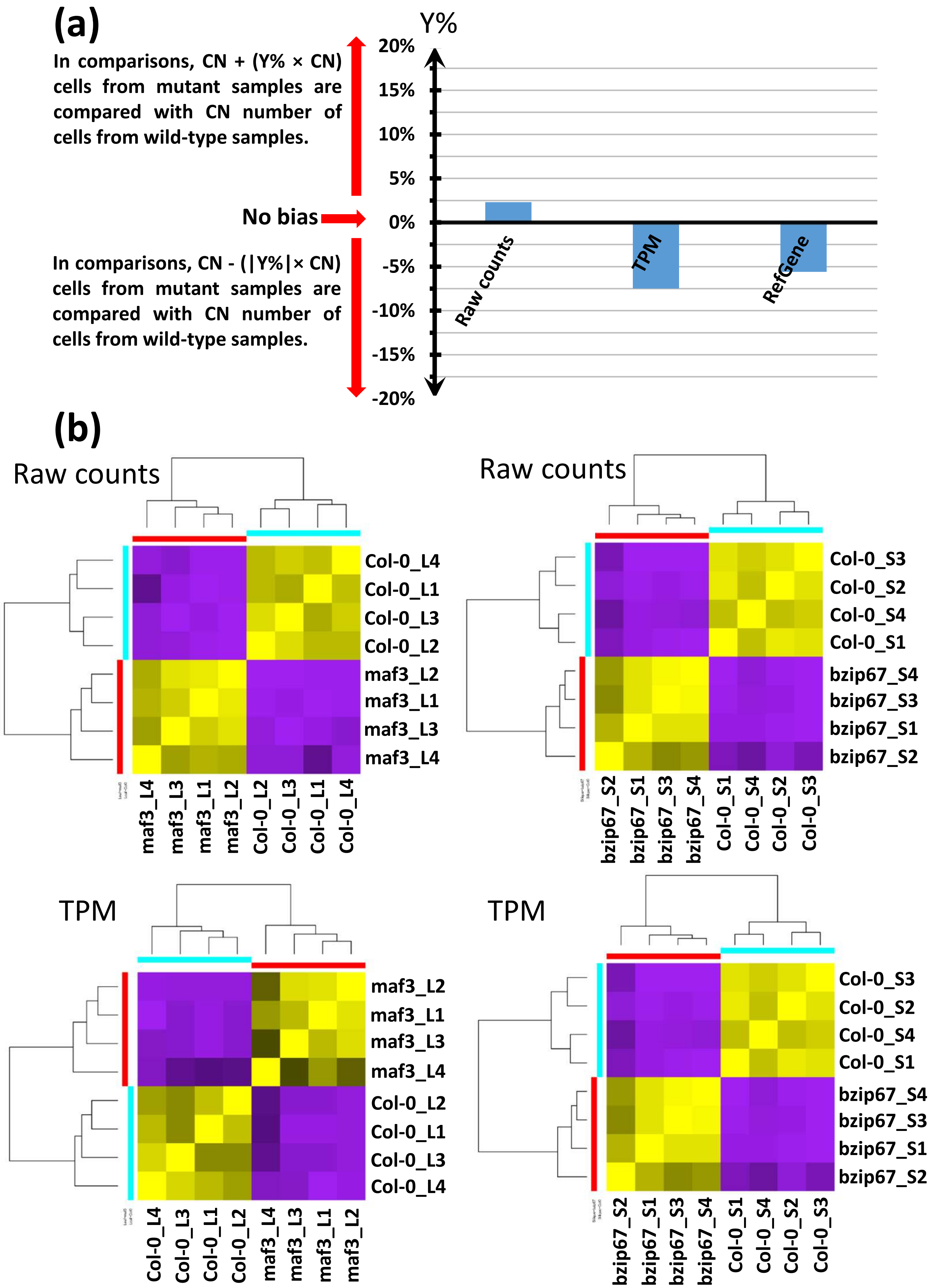


**Fig. S6 continued.**

**(b)**

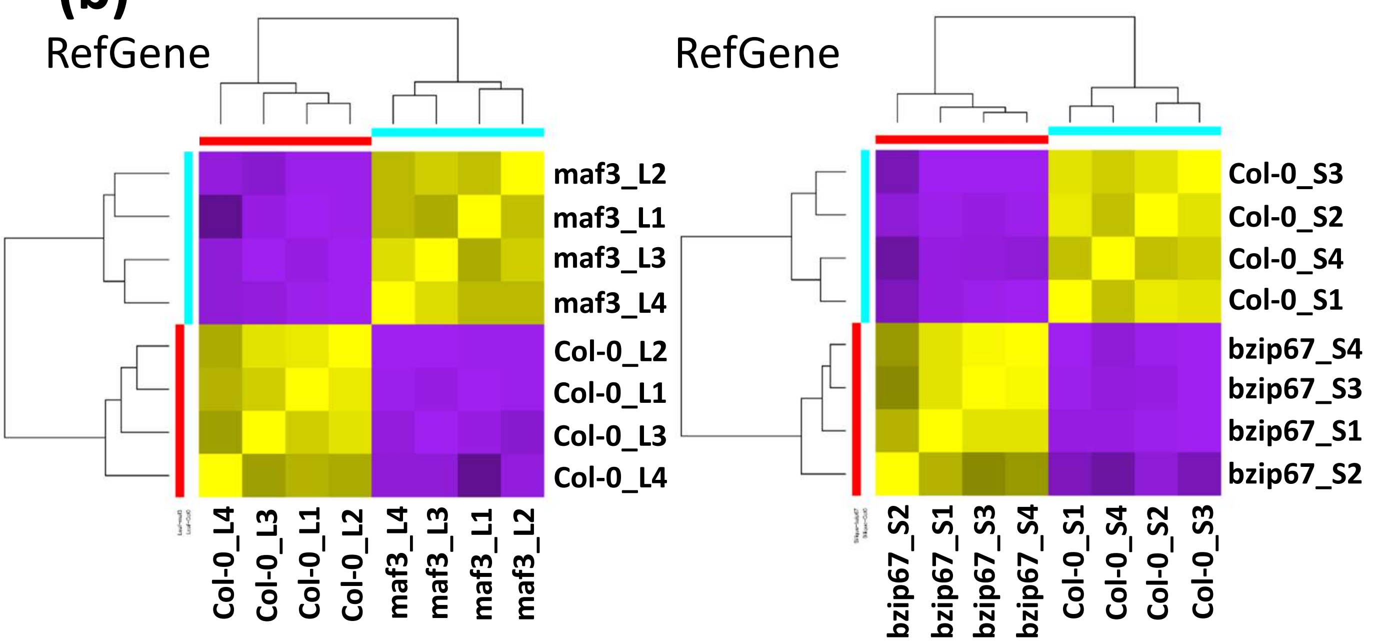


**Fig. S6 Data normalization for *Arabidopsis* transcriptome experiments.** (a) Inter-treatment bias levels for cell-number in *Arabidopsis* samples. The bias for cell-number was examined while comparing the mutant plants with the wild-type plants. CN: cell number. (b) Heatmap of the sample clusters based on the expression levels of differentially expressed (DE) genes. DE genes, i.e., FDR < 0.001 & FC ≥ 2.35 in *bzip*P67 experiment samples and, FDR < 0.001 & ≥ 3.18 in *maf*3 experiment samples. First, DE genes were identified on raw count data using edgeR and afterwards, raw count data, TPM (transcript per million reads) data, and reference gene (GADPH coding gene AT1G13440) based corrected data were used to obtain the Pearson correlation heatmaps. 1-4: replicates, L: leaf, S: silique.

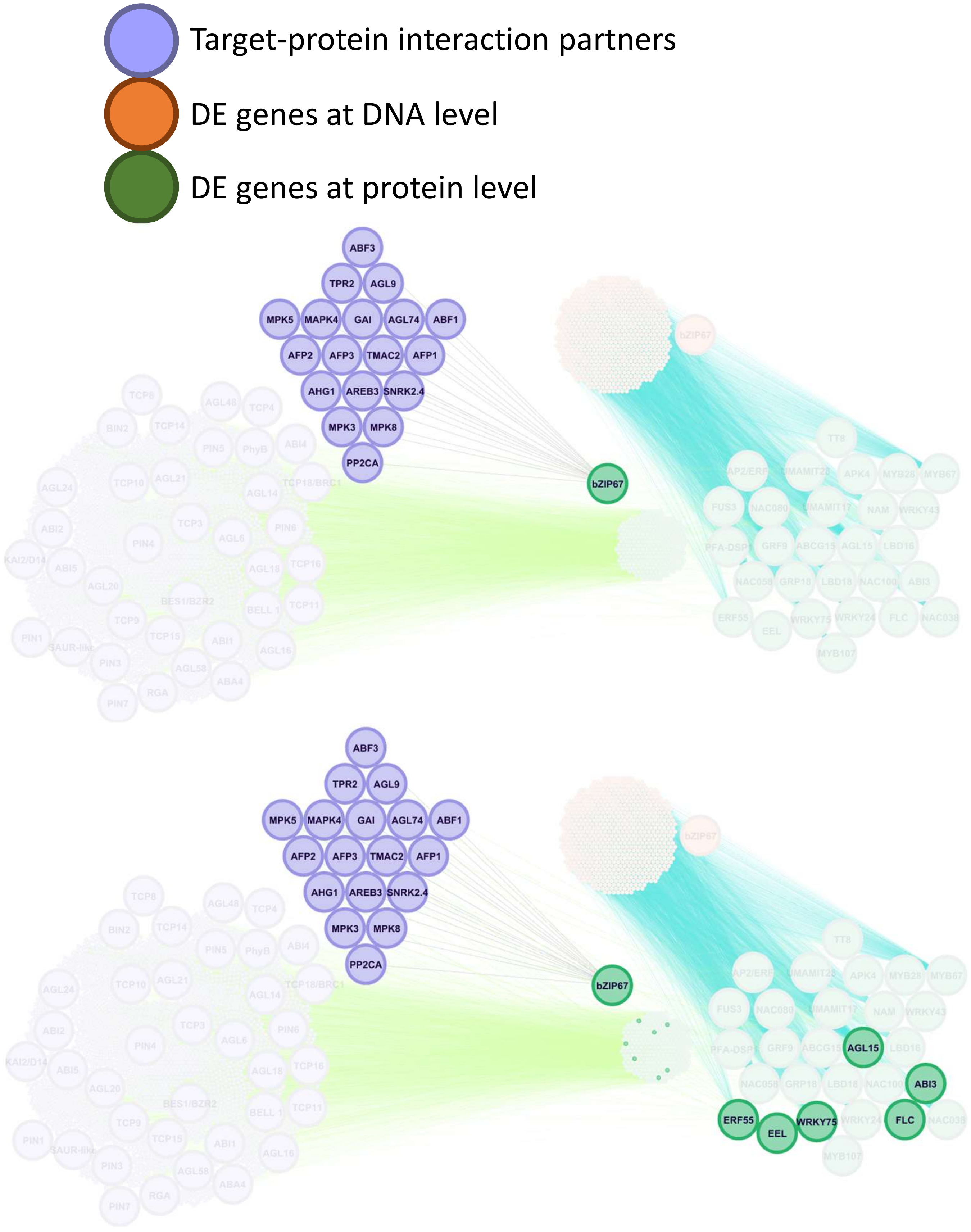


**Fig. S7 continued.**

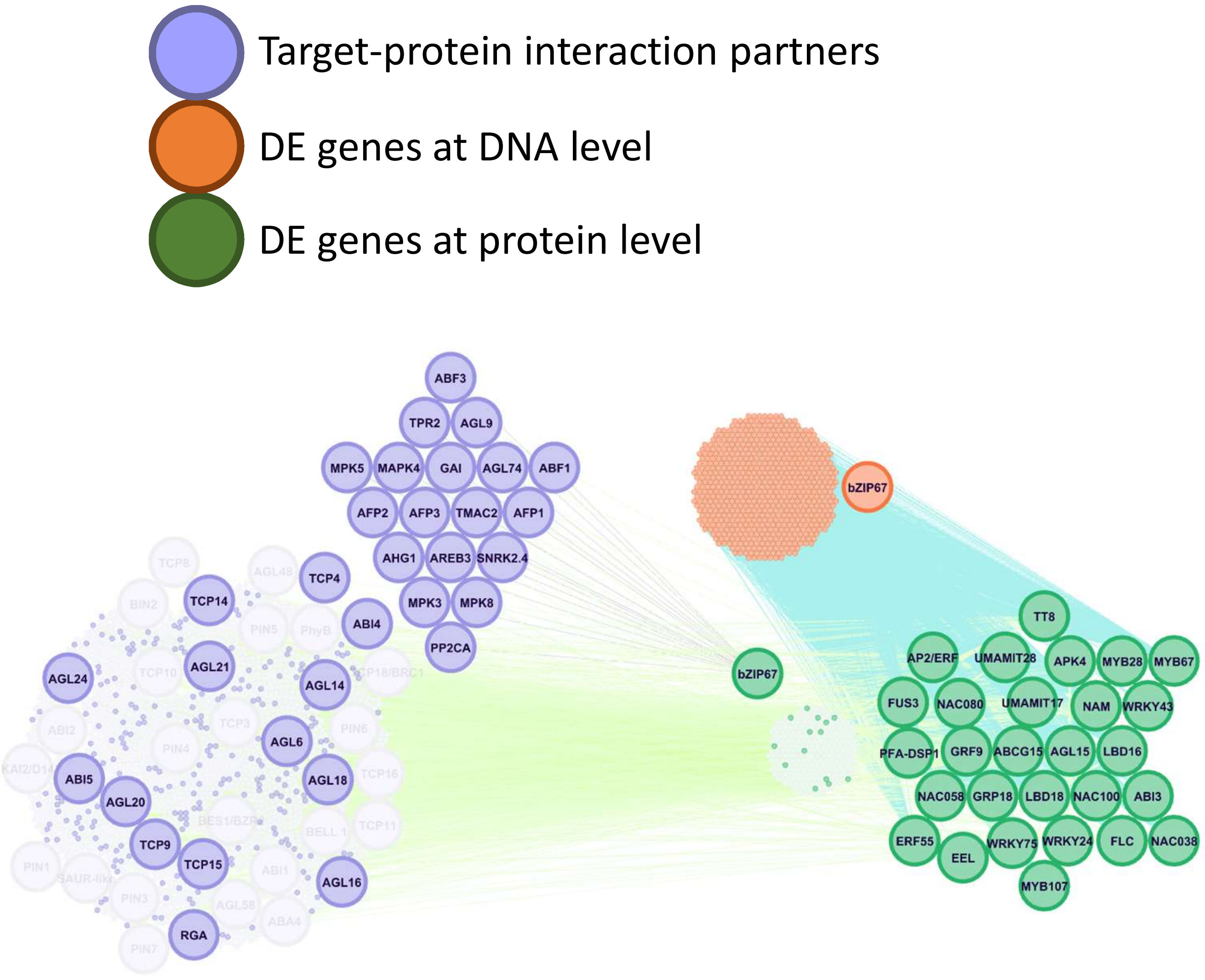


**Fig. S7 Protein-protein and protein-DNA interaction network for the differentially expressed genes and their interacting partners.** Protein-protein and protein-DNA interaction network are shown in a cascade mode with bZIP67 transcription factor as starting point from top panel towards to the second and third, i.e., bottom, panels.

**(a)**

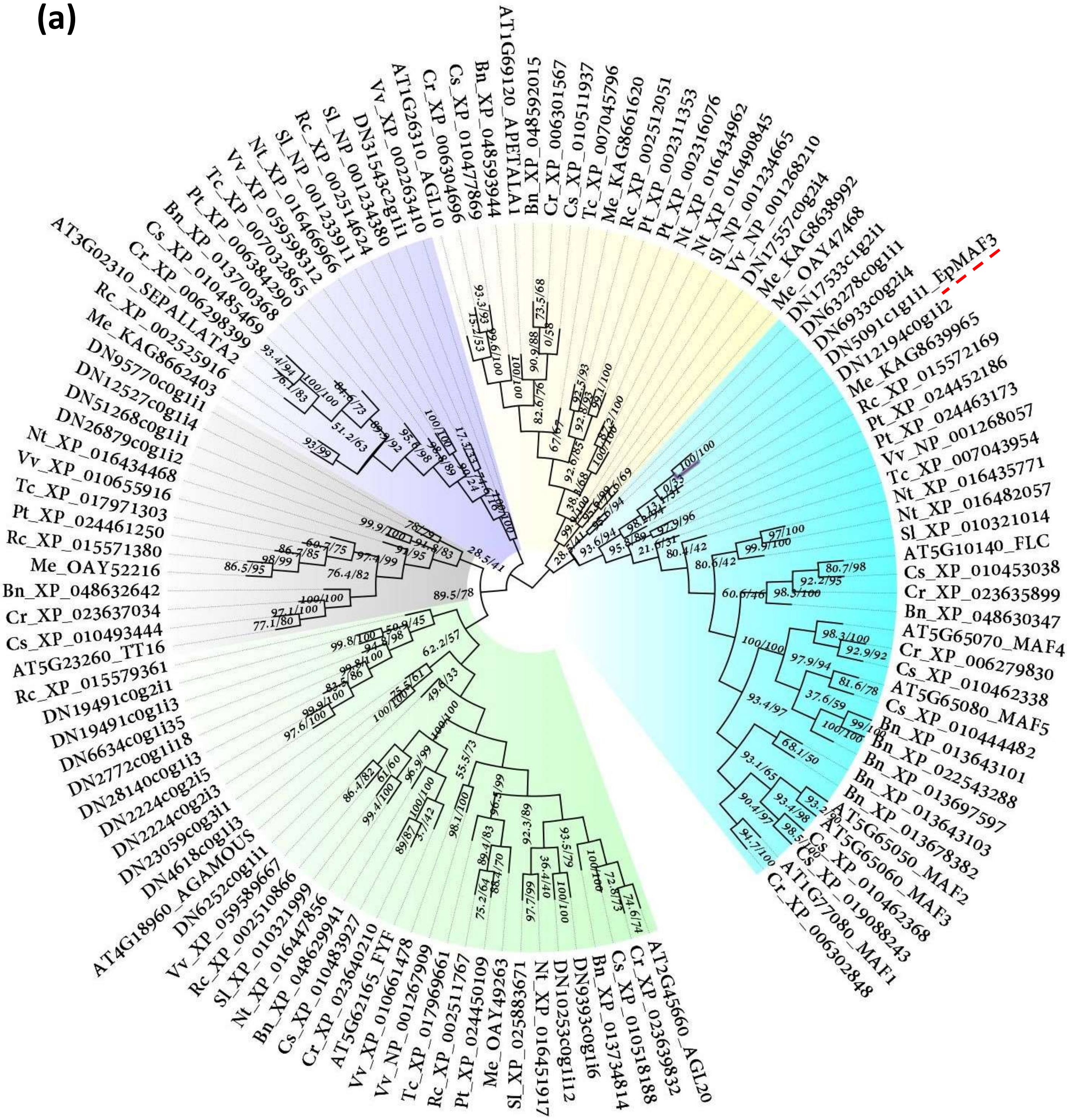


**Fig. S8 continued.**

**(b)**

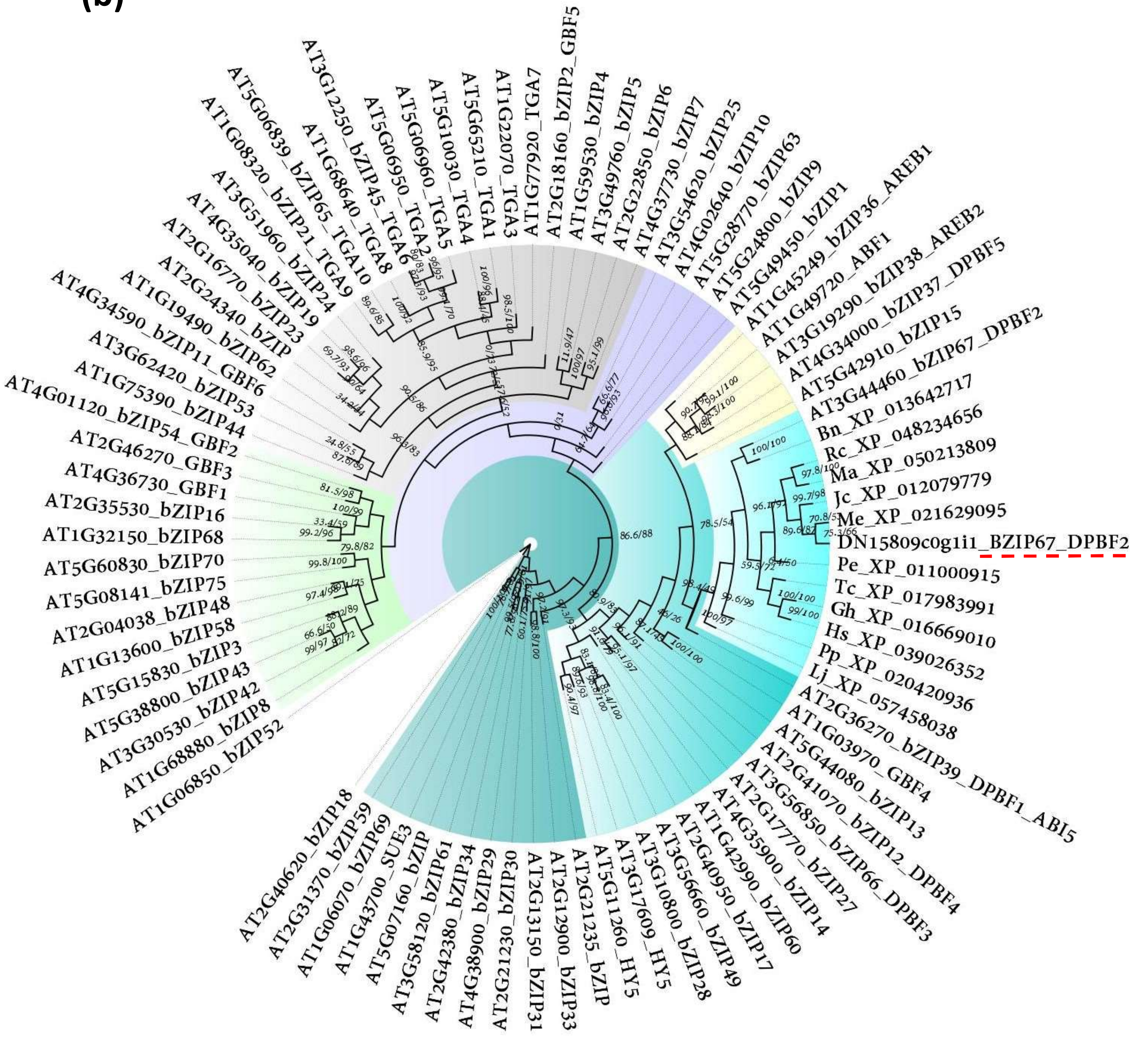


**Fig. S8 continued.**

**Fig. S8 Phylogenetic analysis for the *bZip*67 and *Maf*3 selected candidate genes.** (a) Phylogenetic tree of *Arabidopsis* MAFs and AGAMOUS like proteins. (b) Phylogenetic tree for *Arabidopsis* bZIP transcription factors. The phylogenetic trees in A and B were built using IQ-TREE. Bootstrapping was set to 1000 (aLRT%/UFBoot%). Taxonomic trees are drawn by the Taxallnomy web application. The NCBI or TAIR accession codes are shown for the sequences. The genus and species initials are illustrated as prefixes; Bn: *Brassica napus*, Cr: *Capsella rubella,* Cs: *Camelina sativa*, Gh: *Gossypium hirsutum*, Hs: *Hibiscus syriacus*, Jc: *Jatropha curcas*, Lj: *Lotus japonicus*, Ma: *Mercurialis annua*, Me: *Manihot esculenta*, Nt: *Nicotiana tabacum*, Pe: *Populus euphratica,* Pp: *Prunus persica*, Pt: *Populus trichocarpa*, Rc: *Ricinus communis*, Sl: *Solanum lycopersicum*, Tc: *Theobroma cacao*, & Vv: *Vitis vinifera*. Accession codes starting with DN are from poinsettia. Poinsettia MAF3 and bZIP67 are underlined in red.

**Table S1 Sequencing statistics for poinsettia and *Arabidopsis* samples**

| | | Sample | Paired reads | Paired reads (trimmed) | Concordant mapping rate |
|---|---|---|---|---|---|
| **Poinsettia** | **No phytoplasma infection** | **2nd & 3rd buds no.1** | 42,806,133 | 99.99% | 89.78% |
| | | **2nd & 3rd buds no.2** | 42,051,378 | 100% | 84.69% |
| | | **2nd & 3rd buds no.3** | 46,182,468 | 100% | 88.11% |
| | **Phytoplasma infected** | **2nd & 3rd buds no.1** | 46,023,889 | 99.99% | 92.38% |
| | | **2nd & 3rd buds no.2** | 46,218,450 | 100% | 90.88% |
| | | **2nd & 3rd buds no.3** | 47,248,433 | 100% | 91.07% |
| | **No phytoplasma infection** | **4th bud no.1** | 47,653,396 | 99.99% | 87.98% |
| | | **4th bud no.2** | 46,790,124 | 99.99% | 88.53% |
| | | **4th bud no.3** | 45,001,353 | 100% | 91.20% |
| | **Phytoplasma infected** | **4th bud no.1** | 46,681,626 | 100% | 85.95% |
| | | **4th bud no.2** | 47,185,566 | 100% | 90.06% |
| | | **4th bud no.3** | 43,120,060 | 100% | 84.64% |
| | **Total or Avg.** | | **546,962,876** | **100%** | **88.77%** |
| **Arabidopsis** | ***maf*3 mutant-leaf** | **maf3-L1** | 58,186,285 | 100% | 91.24% |
| | | **maf3-L2** | 58,897,673 | 100% | 90.94% |
| | | **maf3-L3** | 50,643,004 | 100% | 90.61% |
| | | **maf3-L4** | 53,153,918 | 100% | 91.52% |
| | **Col-0 leaf** | **Col-0-L1** | 50,843,500 | 100% | 90.61% |
| | | **Col-0-L2** | 47,251,155 | 100% | 91.22% |
| | | **Col-0-L3** | 63,323,400 | 100% | 90.62 |
| | | **Col-0-L4** | 48,765,525 | 100% | 89.98% |
| | ***bzip*67 mutant-silique** | **bzip67-S1** | 54,826,806 | 100% | 90.37% |
| | | **bzip67-S2** | 49,379,510 | 100% | 92.04% |
| | | **bzip67-S3** | 46,905,396 | 100% | 90.34% |
| | | **bzip67-S4** | 50,048,204 | 100% | 91.59% |
| | **Col-0 silique** | **Col-0-S1** | 51,911,622 | 100% | 91.41% |
| | | **Col-0-S2** | 72,290,576 | 100% | 91.72% |
| | | **Col-0-S3** | 57,634,268 | 100% | 91.32% |
| | | **Col-0-S4** | 68,926,234 | 100% | 86.84% |
| | **Avg.** | | **55,186,692** | **100%** | **91.0%** |

**Table S2 De novo assembly statistics of poinsettia transcriptome**

| | Length (bases) | Number |
|---|---|---|
| **Genes** | 86,137,880 | 127,338 |
| **Transcripts** | 181,636,050 | 213,118 |
| **N50** | 1,398 | 39,381 |
| **N40** | 1,717 | 27,657 |
| **N20** | 2,087 | 18,048 |
| **N20** | 2,585 | 10,191 |
| **N10** | 3,445 | 4,028 |
| **Median** | 502 | |
| **Average** | 847.4 | |
| **Longest** | 16,238 | |
| **Shortest** | 179 | |
